# A new tool for quantifying mitochondrial RNA processing intermediates reveals increased levels of immature mRNA species under oxidative stress

**DOI:** 10.64898/2026.10.30.763231

**Authors:** Ulas Ozkurede, Shreya Rajhans, Federica Accornero, Julia C. Liu

**Author notes:** Lead contact: Julia C. Liu.

## Abstract

Mitochondrial gene expression in mammals requires the processing of polycistronic transcripts into mature RNAs, generating multigene maturation intermediates. Yet these species and their potential association with mitochondrial stress remain poorly characterized.

We analyzed intact RNA species in healthy mouse hearts through direct RNA sequencing and identified polyadenylated mitochondrial transcripts containing complete mRNA sequences with uncleaved tRNA extensions, which we termed immature mRNAs (imRNAs). Guided by these findings, we developed a new computational tool to estimate imRNA levels using conventional short-read RNA-seq datasets. Analysis of mouse hearts deficient in mitochondrial superoxide dismutase 2 (SOD2), a model of mitochondrial oxidative stress, revealed selectively elevated levels of imRNAs retaining 5’ tRNA extensions, without increases in those with 3’ tRNA extensions. Comparative analyses of several mouse models with defects in mitochondrial respiration, transcription, translation, or proteostasis did not reproduce this pattern. A mouse model of mitochondrial import stress with reduced ROS generation exhibited decreased levels of imRNAs retaining 5’ tRNA extensions. Conversely, human induced pluripotent stem cells treated with rotenone, which elevates mitochondrial ROS, showed increased levels of imRNAs retaining 5’ but not 3’ tRNA extensions, reproducing the pattern observed in SOD2-deficient mouse hearts.

These findings identify an imRNA signature potentially associated with mitochondrial oxidative stress. The presence of imRNAs described here suggests that widely used transcriptome analysis pipelines may count these species as mature mRNAs, overestimating mitochondrial mRNA levels and overlooking defects in RNA maturation. By enabling retrospective analysis of processing intermediates, our tool provides access to previously unexamined information in existing RNA-seq datasets.

## 1. Introduction

Mitochondria are the only organelles that contain extranuclear genomes in animal cells. Mitochondrial oxidative phosphorylation (respiration) relies on both nuclear gene expression and the local production of proteins encoded by mitochondrial DNA (mtDNA). Mammalian mtDNA encodes 13 proteins, all essential subunits of the oxidative phosphorylation machinery, as well as two ribosomal RNAs and 22 transfer RNAs that enable the local translation of mitochondrial mRNAs. The remaining proteins required for respiration and mitochondrial gene expression are encoded in the nuclear genome. Mitochondrial protein synthesis is essential for mammalian physiology. Reflecting this, loss of genes required for mitochondrial transcription, RNA processing, or translation is lethal in mice, while partial defects in these processes are associated with several pathologies in humans, including intellectual disability (Olahova et al., 2021), liver failure (Stiles et al., 2016), and cardiomyopathy (Haack et al., 2013).

Current understanding of the regulation of mitochondrial gene expression lags far behind that of nuclear gene expression, especially with regard to posttranscriptional mechanisms. A major factor is that, while nuclear genome editing methods have been developed and successfully employed for decades, molecular tools for editing mtDNA are still lacking. Mitochondrial mRNA maturation differs fundamentally from its nuclear counterpart. The compact mitochondrial genome is transcribed into long polycistronic RNAs containing multiple rRNA, tRNA, and mRNA sequences. Individual RNA species are released through endonucleolytic cleavage, largely at tRNA boundaries, and undergo further maturation. In contrast to the ordered steps of nuclear mRNA maturation (Moore & Proudfoot, 2009), mitochondrial RNA processing does not necessarily proceed sequentially along a precursor transcript. Previous studies suggest that cleavage and polyadenylation at a junction site can, in theory, generate a mature 3’ end with a polyA tail while upstream junctions in the same transcript remain unprocessed. The abundance of these partially processed species could reveal changes in mitochondrial RNA metabolism. However, to our knowledge, no dedicated bioinformatics tool is available to quantify them in high-throughput short-read RNA sequencing data. Reads spanning uncleaved gene boundaries provide information specific to incompletely processed RNA, but standard gene-level quantification does not distinguish these species. This limitation leaves potentially informative features of mitochondrial RNA maturation unexamined in existing short-read RNA-seq datasets.

How mitochondrial RNA maturation responds to oxidative stress is of particular interest, as the electron transport chain (ETC) in the mitochondrial inner membrane is a major source of reactive oxygen species (ROS). Superoxide dismutase 2 (SOD2), located in the mitochondrial matrix, limits superoxide levels by catalyzing its conversion to hydrogen peroxide. Accordingly, loss of SOD2 causes mitochondrial ROS (mtROS) elevation and oxidative stress. While mice lacking the cytosolic superoxide dismutase SOD1 (Ho et al., 1998) or the extracellular enzyme SOD3 (Carlsson, Jonsson, Edlund, & Marklund, 1995) can reach adulthood, those without SOD2 die within two weeks of birth (Lebovitz et al., 1996; Li et al., 1995), underscoring the importance of mtROS clearance. Unexpectedly, mice lacking SOD2 in muscle tissues, including the heart, can survive development and reproduce before they develop dilated cardiomyopathy in early adulthood. This suggests additional mechanisms that allow sustained mitochondrial respiration under conditions of compromised mtROS clearance. Historically, mtROS have been thought to cause mutations in the mitochondrial genome, contributing to disease and aging (Miquel, Economos, Fleming, & Johnson, 1980; Richter, Park, & Ames, 1988; Schriner et al., 2005). However, diligent work by the Larsson group suggested that increased mtROS does not necessarily produce the anticipated damage to mitochondrial nucleic acids. In muscle-specific SOD2-deficient mice, increased oxidative stress was not accompanied by increased mutation rates in mtDNA or mitochondrial RNA (Kauppila et al., 2018). Whether mitochondrial RNA maturation is altered under these conditions remains unclear.

We combined Nanopore direct RNA sequencing of healthy mouse hearts with computational analysis of publicly available short-read RNA-seq datasets to investigate mitochondrial mRNA maturation under oxidative stress. Direct RNA sequencing identified polyadenylated transcripts containing complete mRNA coding sequences with uncleaved tRNA extensions, which we termed immature mRNAs (imRNAs). We developed a computational tool to estimate their relative abundance from reads spanning mitochondrial tRNA–mRNA and mRNA–tRNA boundaries, distinguishing imRNAs retaining 5’ tRNA extensions from those retaining 3’ tRNA extensions. Applying this approach revealed a distinct imRNA signature in SOD2-deficient mouse hearts. Comparisons with other mitochondrial perturbations and rotenone-treated human cells suggested an association between this imRNA pattern and mitochondrial oxidative stress. Our findings reveal a feature of mitochondrial RNA metabolism that is obscured by conventional gene-level analysis and provide a new approach for investigating it in existing RNA-seq datasets.

## 2. Methods

### 2.1. RNA-seq datasets from previously published models

#### *Sod2*-mKO mice

*Sod2^flox/flox^* mice (Strassburger et al., 2005) harbor a modified *Sod2* gene with a *loxP*-flanked region encompassing the third exon, removal of which results in an inactive SOD2 protein (Li et al., 1995). *Ckmm-cre* transgenic mice express Cre recombinase under the control of the muscle creatine kinase (*Ckmm*) promoter (J. Wang et al., 1999). The double-transgenic *Sod2^fl/fl^*;*Ckmm-cre* (heart and skeletal muscle-specific *Sod2* knockout; *Sod2*-mKO) and *Sod2^fl/fl^* (control) mice have been generated and studied by the Larsson group (Kauppila et al., 2018). Publicly available raw RNA sequencing files (NCBI BioProject PRJEB25258; Illumina, single-end, 150-bp, ∼35M reads/sample) from three pairs of 10–11-week-old control and *Sod2*-mKO mice (Supplementary Table 5) were utilized in transcriptome, mitochondrial mRNA expression, annotation-agnostic mtDNA expression, and mitochondrial immature mRNA (imRNA) analyses presented in this study.

#### *Ant1*-Tg mice

*Ant1^Tg^*/+ mice (X. Wang, Middleton, Tawil, & Chen, 2022) have extra copies of the *Ant1* gene with its native promoter sequence, resulting in *Ant1* overexpression. The Chen group recently reported RNA-seq analyses of quadriceps skeletal muscle tissue samples from these mice (Brennan et al., 2026). Raw RNA sequencing files (NCBI BioProject PRJNA1188003; Illumina NextSeq 2000, paired-end, 2 × 100-bp, ∼25M reads/sample) from four pairs of 2-month-old wild-type and *Ant1*-Tg animals (Supplementary Table 5) were utilized in mitochondrial mRNA expression and imRNA analyses reported here.

#### *Mterf4*-mKO, *Tfam*-mKO, and *Polrmt*-mKO mice

*Mterf4^LoxP/LoxP^* (Cámara et al., 2011), *Tfam^LoxP/LoxP^* (Larsson et al., 1998), and *Polrmt^LoxP/LoxP^* mice (Kühl et al., 2016) have, respectively, modified *Mterf4*, *Tfam*, and *Polrmt* genes with *loxP*-flanked regions, excision of which results in deficiencies in the corresponding proteins. Crosses of these strains with mice expressing Cre under the control of the *Ckmm* promoter (MGI:2177173; Tg[Ckmm-cre]1Lrsn) yield the muscle-specific knockout mouse models *Mterf4^LoxP/LoxP^*; +/*Ckmm-cre* (*Mterf4*-mKO) (Cámara et al., 2011), *Tfam^LoxP/LoxP^*; +/*Ckmm-cre* (*Tfam*-mKO) (Hansson et al., 2004), and *Polrmt^LoxP/LoxP^*; +/*Ckmm-cre* (*Polrmt*-mKO) (Kühl et al., 2014). These models have been employed in RNA-seq experiments for comparative analyses (Kühl et al., 2017). In the current study, raw RNA sequencing datasets (NCBI BioProject, PRJNA378878; Illumina HiSeq 4000, paired-end, 2 × 75-bp, ∼50M reads/sample) generated from heart tissues of *Mterf4*-mKO, *Tfam*-mKO, and *Polrmt*-mKO mice and corresponding controls (Supplementary Table 5) were utilized in mitochondrial mRNA expression and imRNA analyses.

#### *Lonp1*-smKO mice

Mice with skeletal muscle-specific deletion of the gene encoding Lon peptidase 1, mitochondrial (*Lonp1*-smKO mice) were previously described and analyzed through RNA sequencing by the Gan group (Xu et al., 2022). In the current study, raw sequencing data (NCBI BioProject PRJNA699182; Illumina HiSeq 4000, paired-end, 2 × 150-bp) generated from gastrocnemius muscle tissues of *Lonp1*-smKO (n = 2) and wild-type (n = 2) mice were utilized in mRNA expression and imRNA analyses.

#### Human iPSCs treated with rotenone

The Sone group exposed human iPS cells (HPS0003) to increasing concentrations of dimethyl sulfoxide (DMSO; 0.001%, 0.01%, 0.1%) or rotenone (0.3 μM, 1 μM, 25 μM) for 48 hours and performed RNA sequencing (Hongen, Sakai, Ito, Qin, & Sone, 2024). Publicly available raw RNA sequencing data obtained from these samples (NCBI BioProject PRJNA1188018; Illumina NovaSeq 6000, paired-end, 2 × 150-bp, ∼30M reads/sample) (Supplementary Table 6) were utilized in mitochondrial mRNA expression and imRNA analyses.

#### Mouse embryonic fibroblasts devoid of mtDNA

MEFs lacking mtDNA (ρ^0^ MEFs), generated from control MEFs with intact mtDNA through ditercalinium dichloride dihydrochloride treatment, were previously utilized in an RNA-seq study by the Teitell group (Shimada et al., 2018). A subset of the publicly available dataset (GEO, GSE111668; SRA, SRP134850) including raw sequencing reads from three control and three ρ^0^ MEF cultures was analyzed in this study to assess the specificity of mouse uncleaved junction identifiers (UJIs) (Supplementary Table 6).

#### Human colorectal carcinoma HCT116 cells lacking mtDNA

HCT116 cells depleted of mtDNA (ρ^0^ HCT cells), generated through ethidium bromide-mediated mtDNA elimination, were analyzed through RNA-seq experiments together with control HCT116 cells containing intact mtDNA by the Savkovic group (Heller et al., 2017). Part of the published RNA-seq dataset (SRA, SRP093357) including raw sequencing reads from three control and three ρ^0^ HCT cultures (Supplementary Table 6) was analyzed in the current study to assess human UJI specificity.

### 2.2. RNA-seq gene-level expression and pathway analysis

For gene-level differential expression analysis, raw RNA-seq data from control and experimental samples were processed using the CHURP bulk RNA-seq pipeline (version 1.1.0) at the Minnesota Supercomputing Institute, University of Minnesota. Briefly, sequencing adapters and low-quality bases were removed using Trimmomatic, with leading and trailing base quality thresholds of 3, a sliding window of 4 bases with a minimum average Phred quality score of 15, and a minimum retained read length of 18 bases. Trimmed reads were aligned to the mouse reference genome (GRCm39, Ensembl release 113) using HISAT2. Gene-level read counts were generated using the featureCounts tool with the corresponding Ensembl annotation (Mus_musculus.GRCm39.113.gtf). Raw counts were imported into the edgeR package in R. Differential expression testing was performed with a quasi-likelihood test in edgeR between two groups. A 0.05 false discovery rate (FDR) correction was applied to the tests.

To identify potential biological processes that are disproportionately represented among downregulated genes, a list of genes with counts per million reads (CPM) > 10 and fold-change < 0.8 was provided as input for the Gene Ontology (GO) Biological Process overrepresentation analysis using the PANTHER tool available on geneontology.org (Ashburner et al., 2000; Consortium, 2026). Normalized gene counts (CPM) of mammalian genes encoding respiratory chain subunits based on MitoCarta3.0 (Pagliarini et al., 2008; Rath et al., 2021) were extracted from the RNA-seq data and presented using a heat map.

### 2.3. Mitochondrial mRNA expression analysis with custom annotations

Reference mitochondrial genome annotation files were downloaded from Ensembl (release 113) for mouse (Mus_musculus.GRCm39.113.gtf) and human (Homo_sapiens.GRCh38.113.gtf). Each file includes 13 protein-coding genes, while mammalian mitochondria produce 11 mature mRNA species. Nine of these mRNAs are monocistronic, while the remaining two are bicistronic, each containing two partially overlapping open reading frames. The first bicistronic mRNA is translated into ATP8 and ATP6; the second into ND4L and ND4 (**fig 1C**). To account for these bicistronic transcripts in mitochondrial mRNA quantification, custom mitochondrial genome annotation files were generated. Briefly, the mouse mRNA annotations for *mt-Atp8* (7766–7969) and *mt-Atp6* (7927–8607) were replaced with a single annotation designated *mt-Atp8–6* (7766–8607), spanning both coding sequences. Similarly, *mt-Nd4l* (9877–10173) and *mt-Nd4* (10167–11544) were combined into a single annotation designated *mt-Nd4l–4* (9877–11544). Corresponding modifications were made to the human mitochondrial genome annotation. Raw RNA-seq reads were aligned to the respective reference mitochondrial genomes using STAR (Dobin et al., 2013). Gene-level read counts for heavy-strand mRNAs were obtained using featureCounts (Liao, Smyth, & Shi, 2014) using the custom mitochondrial annotations described above. Mitochondrial mRNA abundance was calculated as transcripts per million reads (TPM).

### 2.4. Annotation-agnostic analysis of mtDNA expression

The mouse mitochondrial genome was divided into 30-nt intervals, generating 543 nonoverlapping sequences covering the first 16,290 nucleotides and one additional interval spanning the last nine nucleotides and the first 21 nucleotides of the circular genome. The abundance of each interval was quantified using a custom Python script by counting RNA-seq reads that contain exact sequence matches. The results were expressed as counts per 100 million mitochondrial reads. Intervals showing differences in abundance with p < 0.05 and |log_2_^R^| > 0.7, where R is the ratio of mean values, were considered differentially expressed.

### 2.5. Wild-type mice

Three-month-old male C57BL/6N mice (n = 3) were used for Nanopore direct RNA sequencing experiments. Mice came from an in-house colony maintained on a C57BL/6N background and carrying a Cre recombinase transgene driven by the cardiomyocyte-specific α-myosin heavy chain promoter (αMHC-Cre; JAX stock no. 011038). The colony was housed in a controlled environment at 22 °C and 40–60% humidity on a 12-hour light/dark cycle, with *ad libitum* access to standard chow diet and water. All animal experiments were approved by the Institutional Animal Care and Use Committee at Brown University (protocol number 23-05-0005).

### 2.6. Direct RNA sequencing

Mice were anesthetized with 2% isoflurane and euthanized through cervical dislocation. Hearts were harvested, snap-frozen in liquid nitrogen, and stored at −80 °C until RNA extraction. Total RNA was extracted from frozen hearts using TRIzol reagent (Invitrogen, Cat. #15596018) after bead homogenization. RNA concentration was measured with the Qubit RNA Broad Range Assay (Thermo Fisher Scientific), and RNA integrity was assessed on an Agilent TapeStation using the High Sensitivity RNA ScreenTape assay; RINe values ranged from 8.1 to 8.8 across all samples.

Direct RNA sequencing libraries were prepared using the Direct RNA Sequencing Kit (SQK-RNA004; Oxford Nanopore Technologies) according to the manufacturer’s protocol. Briefly, 1 µg of total RNA per sample was ligated to the oligo(dT)-containing RT Adapter, which selectively captures polyadenylated RNAs, and a complementary cDNA strand was synthesized using Induro Reverse Transcriptase (New England Biolabs). The RNA–cDNA hybrids were purified with Agencourt RNAClean XP beads (Beckman Coulter) and ligated to the RNA Ligation Adapter. After a second bead cleanup, final libraries were quantified with the Qubit dsDNA HS Assay (Thermo Fisher Scientific).

Each library was loaded onto a PromethION RNA flow cell (FLO-PRO004RA; Oxford Nanopore Technologies) and sequenced using MinKNOW (v25.09.16). Basecalling was performed in real time with Dorado v1.1.1 (basecalling server v7.11.2) using the super-accuracy (SUP) RNA004 model (130 bps). Reads with a mean quality score ≥ 10 were classified as pass reads and used for downstream analysis.

### 2.7. Characterization of mtRNA species in direct RNA sequencing data

Nanopore direct RNA sequencing reads were filtered to retain those representing mitochondrial transcripts. Each mitochondrial read was analyzed using a 30-nt sliding window to generate an ordered series of subsequences. These subsequences were matched to the reference mitochondrial genome, requiring perfect 30-nt sequence matches, to identify the corresponding genes and generate an ordered series of gene identifiers. Matches to noncoding regions and partial matches to two neighboring genes were removed, while consecutive occurrences of the same gene identifier were collapsed into a single entry, generating a simplified representation of the genes covered by each transcript while preserving their original order. These ordered series were subsequently converted into gene-type-level transcript topologies by replacing individual gene identifiers with their corresponding RNA types. For example, the original series [*Nd1*, *Nd1*, *Nd1*, *Nd1–TrnI*, *TrnI*, *TrnI*, *TrnI–Nd2*, *Nd2*, *Nd2*] was reduced to [*Nd1*, *TrnI*, *Nd2*] and subsequently converted to [mRNA, tRNA, mRNA]. The last representation described the sequential arrangement of RNA types within each transcript.

Using the gene-type-level transcript topology data, transcripts were classified based on their RNA composition: rRNA only, tRNA only, mRNA only, rRNA and tRNA, mRNA and tRNA, or all three RNA types. The percentage of transcripts in each class was calculated and reported for individual mice. Transcripts containing both mRNA and tRNA sequences were further classified according to the number of distinct mRNA, tRNA, and rRNA species present in each transcript. For example, the gene-type-level transcript topology [mRNA, tRNA, mRNA] corresponded to the class (2 mRNA, 1 tRNA, 0 rRNA). Classes represented by at least 100 reads were reported.

### 2.8. Development of a new tool for imRNA analysis using short-read RNA-seq data

#### Immature mRNAs

An immature mRNA (imRNA) was defined as an RNA species that contains, in addition to the complete coding sequence present in a mature mRNA and a 3’ polyA tail, an unremoved 5’-end or 3’-end extension retained due to incomplete processing (**fig 4B**).

#### Uncleaved junction identifiers

An uncleaved junction (UJ) identifier (UJI) was defined as a 30-nt sequence spanning a boundary between two mitochondrial genes. The presence of mitochondrial imRNA species were inferred by read sequences that contain UJIs spanning tRNA–mRNA or mRNA–tRNA junctions.

#### UJI selection

Based on the distribution of potential 30-nt sequences between two neighboring genes, up to 29 potential UJIs could be defined for each junction, the exact number depending on the length of the intergenic sequence separating the consecutive genes. RNA-seq reads often lack the 5’ ends of isolated transcripts, resulting in a lower coverage of 5’ sequences. This effect, known as 5’-end truncation or 5’-end underrepresentation depending on the sequencing approach, has been noted in both short-read (Adiconis et al., 2013) and direct (Workman et al., 2019) RNA sequencing. We found that minimizing the portion of the UJI derived from the upstream gene improved UJ detection. This observation guided our selection of candidate UJIs, which were then validated for specific detection of mtDNA-encoded transcripts. The sequences of the final sets of mouse and human UJIs are provided in Supplementary Tables 1 and 2.

#### Analysis of nuclear-encoded mitochondrial DNA segments for specific UJI detection

Nuclear genomes contain numerous sequences with high similarity to mtDNA, known as nuclear-encoded mtDNA segments (NUMTs). To be able to use the UJIs as indicators of mitochondrial imRNAs, we had to confirm that they can specifically detect mitochondrial transcripts and not NUMT-derived nuclear transcripts.

To generate complete lists of NUMTs present in mouse (NCBI RefSeq assembly, GCF_000001635.27) and human (NCBI RefSeq assembly, GCF_000001405.40) genomes, we blasted the mtDNA sequences against nuclear genomes.

The following RNA-seq coverage data from mouse tissues were visualized on NCBI Genome Data Viewer: heart (Mus musculus, adult-8wks, SAMN00849390, filtered), kidney (Mus musculus, adult-8wks, SAMN00849385, filtered), liver (Mus musculus, adult-8wks, SAMN00849386, filtered), skeletal muscle (Mus musculus, SAMN01766820, filtered), spleen (Mus musculus, adult-8wks, SAMN00849388, filtered), and whole brain (Mus musculus, E14.5, SAMN01164141, filtered).

#### Biological validation of UJI specificity using ρ^0^ cells

The specificity of uncleaved junction identifiers (UJIs) for mitochondrial transcripts was evaluated using RNA-seq data from control cells with intact mtDNA expression and cells devoid of mtDNA (ρ^0^ cells), which were derived from the control cells. Control cells were used as positive controls for UJI detection in mitochondrial transcripts, whereas ρ^0^ cells were used to assess false-positive matches potentially arising from nuclear mitochondrial DNA segments (NUMTs). Mouse UJIs were evaluated in control and ρ^0^ mouse embryonic fibroblasts (MEFs), and human UJIs in control and ρ^0^ HCT116 cells (Supplementary Table 6). Reads containing UJIs were counted using a Python script. A read was considered a match only if it included a continuous 30-nt interval identical to that of the specified UJI. Counts were normalized to the number of reads analyzed and expressed as counts per billion reads (CPB).

#### Quantitation of imRNA levels with normalization

Reads containing UJIs were counted using a Python script, requiring an exactly matching 30-nt continuous sequence. In each study, UJIs consistently identified in at least 10 reads per biological replicate were considered detected and utilized in subsequent imRNA quantification. The relative abundance of corresponding imRNA species was calculated using Equations (1) and (2):

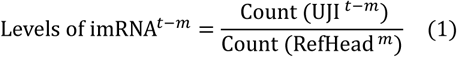

In Equation (1), imRNA*^t–m^* denotes an imRNA species of type I (imRNA with 5’ tRNA) containing an uncleaved junction between the upstream tRNA *t* and downstream mRNA *m*; UJI*^t–m^* denotes the uncleaved junction identifier corresponding to this junction; and RefHead*^m^* denotes a reference sequence within mRNA *m*, located at a predefined distance *d* downstream of its 5’ end.

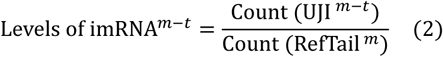

In Equation (2), imRNA*^m–t^* denotes an imRNA species of type II (imRNA with 3’ tRNA) containing an uncleaved junction between the upstream mRNA *m* and downstream tRNA *t*; UJI*^m–t^* denotes the uncleaved junction identifier corresponding to this junction; and RefTail*^m^* denotes a reference sequence within mRNA *m*, located at a predefined distance *d* upstream of its 3’ end.

In Equations (1) and (2), the function *Count* (*x*) returns the number of reads satisfying the exact match requirements for the 30-nt sequence *x*. The distance *d* is identical for both imRNA types, with a default value of 100 nt.

As an additional quality check, the pipeline also quantifies each imRNA species relative to a reference region at the center of its cognate mRNA. In the analyses presented here, normalization to this central reference region yielded no qualitative differences compared with normalization using Equations (1) and (2), which account for potential intragenic expression differences. Discrepancies between the two normalization approaches in a particular study may indicate an mtDNA variant within either reference region.

Experimental model-specific mtDNA mutations within reference or UJI-targeted regions may interfere with exact sequence matching. For variants within a reference region, *d* can be adjusted to shift the reference window to a mutation-free region. The preferred approach, applicable to variants in either region, is to use a custom mtDNA reference sequence incorporating the model-specific variants. In this study, in addition to standard human (NC_012920.1) and mouse (NC_005089.1) mtDNA reference sequences, custom mtDNA sequences with known strain-specific variants were used for imRNA analysis of datasets from mice with different genetic backgrounds.

The Python code and associated resources will be deposited in an online repository upon publication.

### 2.9. Analysis of imRNAs in mitochondrial stress models

Levels of imRNA species were estimated using the imRNA analysis tool described above. The tool takes as input, in addition to raw sequencing data, three reference files: mitochondrial genome (mtDNA.fasta), an annotation file (mtDNA_map.json) describing gene positions, and a UJI file (mtDNA_UJI.json) describing UJIs and their positions. The actual nucleotide sequences for UJIs and reference mRNA sections are extracted from the genome file based on the positions defined in the UJI file. Prior to analysis, each dataset was manually screened for potential mtDNA sequence variants at UJI sites specific to the mouse colony or cell line. Based on detected variations and reported genetic backgrounds, either available strain-specific genomes or custom genomes incorporating the variants were used. For the *Lonp1*-mKO dataset, the C57BL/6J mitochondrial reference genome was used. For datasets from *Sod2-mKO*, *Mterf4*-mKO, *Tfam*-mKO, and *Polrmt*-mKO, and *Ant1*-overexpressing mice, a custom mitochondrial genome incorporating three variants relative to the C57BL/6J reference was used. This genome included two single-nucleotide substitutions, T4891C and T9461C, and an adenine insertion within a tract of eight consecutive adenines beginning at position 9821. Gene annotations and UJI definitions were adjusted accordingly, with all annotation coordinates downstream of the insertion increased by one nucleotide. Of the three variants, only T9461C altered a UJI sequence, introducing a single-nucleotide substitution in UJI *G–Nd3* (supplementary table 1), which was automatically updated by the pipeline based on the reference genome provided. In the analysis of the human iPSC dataset, the parameter *d*, used to assign reference mRNA sequences for normalization, was changed from the default value of 100 to 200 to avoid an mtDNA sequence variation identified within a reference intergenic sequence.

Estimated imRNA levels were reported for individual samples, with no exclusions. For comparisons between groups, t-test p values were corrected for multiple comparisons, and corrected p values below 0.05 were reported on the graphs. Corrected p values below 0.001 were reported as p < 0.001 rather than exact values. Statistical analyses were not performed for the human iPSC dataset, as each culture condition was represented by a single replicate, nor for the *Lonp1*-smKO dataset, as only two mice per group were included in the study.

## 3. Results

### 3.1. Mito-nuclear discordance in respiratory chain expression in SOD2-deficient hearts

We analyzed the *Sod2*-mKO RNA-seq data generated by the Larsson group (NCBI BioProject PRJEB25258; see Methods) and detected a robust downregulation of nuclear-encoded respiratory chain genes in SOD2-deficient mouse hearts (GO term “oxidative phosphorylation”; 9.61-fold enrichment; p = 8.98 × 10^-37^; FDR = 2.19 × 10^-33^). Examination of all nuclear-encoded subunits of the mammalian ETC (Pagliarini et al., 2008; Rath et al., 2021) revealed that, of the 59 differentially expressed subunits, 57 were downregulated, while the only known inhibitory subunit, *Atp5if1*, was upregulated (**fig 1A, S1**). These data indicated a coordinated change in nuclear gene expression consistent with suppression of ETC machinery in cardiac muscle lacking an intact *Sod2* gene. Of note, one gene, *Ndufa11*, deviated from this pattern.

In contrast to nuclear-encoded ETC subunits, none of the 13 mtDNA-encoded subunits showed significant differential expression in SOD2-deficient hearts (**fig 1B, S1**). As noted by others (Jiang et al., 2019; Torres-Gonzalez & Makova, 2022; Vandiver, Herbst, Stothard, & Wanagat, 2025), the reference genome annotations commonly used for gene expression analyses are not optimal for quantifying mitochondrial mRNAs, warranting the use of custom mtDNA annotations (see Methods). We therefore performed an additional analysis of mitochondrial mRNA expression using a custom mtDNA annotation that defines 11 mRNA loci rather than assigning the 13 mtDNA-encoded open reading frames as separate mRNAs (**fig 1C**). This approach reduced the intragroup variations in the estimated levels of *Atp8* and *Atp6* mRNA observed with the standard annotation, suggesting that these were annotation-related artifacts. However, we still found no significant decreases in total (**fig 1D**), ETC complex-specific (**fig 1E**), or individual levels of the 11 mitochondrial mRNAs (**fig 1F**).

The apparent lack of detectible changes in mitochondrial RNA abundance, combined with previous findings indicating no increase in mtDNA or mtRNA mutation rates (Kauppila et al., 2018), suggested that mtROS might not induce detectable effects on the integrity or levels of mitochondrial transcripts. Our observations also implied a marked discrepancy between nuclear (**fig A**) and mitochondrial (**fig 1B, F**) expression of respiratory chain genes in SOD2-deficient hearts. These prompted us to investigate mitochondrial transcripts further using alternative approaches to determine whether changes not captured by conventional analyses were present.

**Figure 1.**
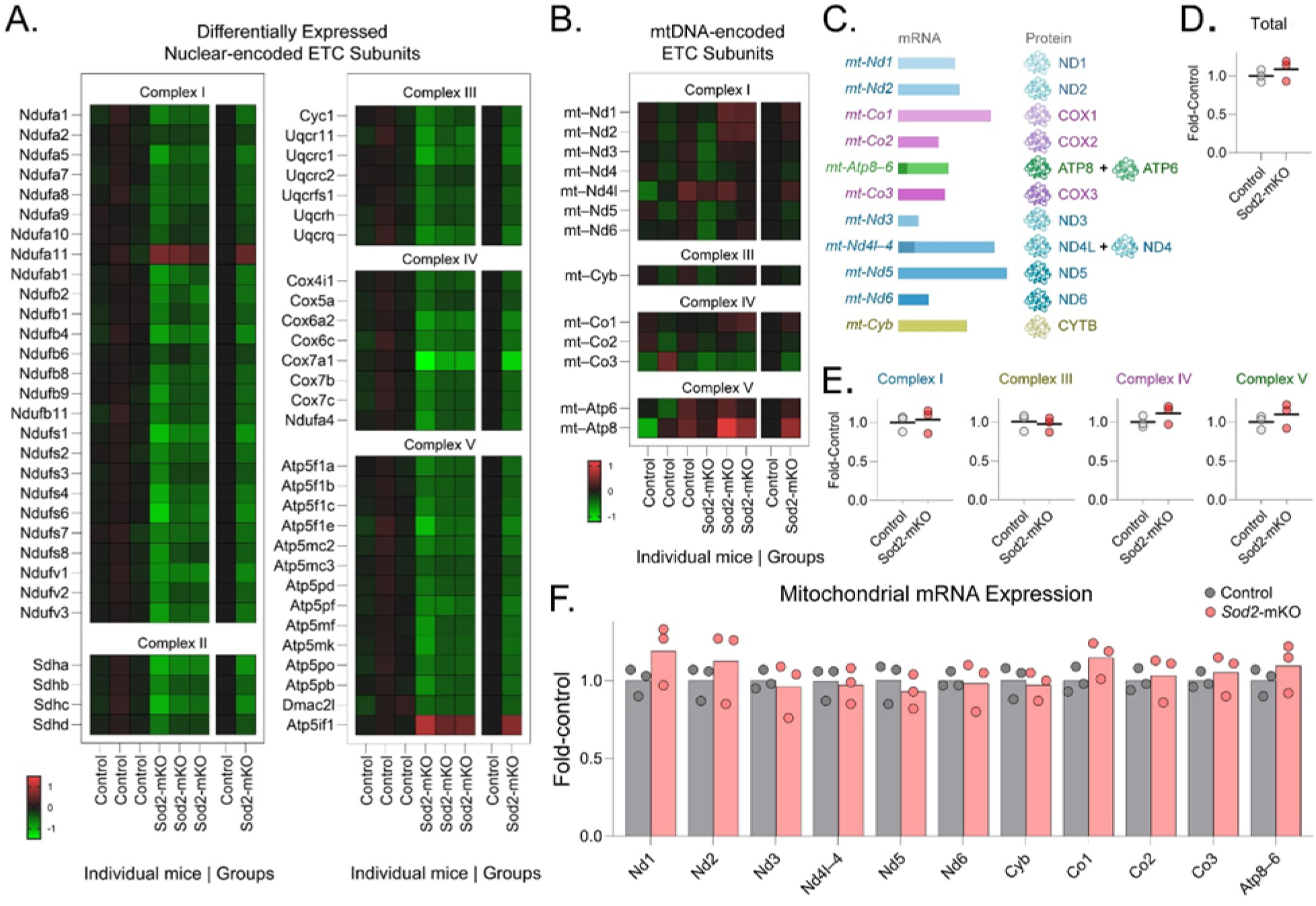
Nuclear and mitochondrial expression levels of respiratory chain subunits in SOD2-deficient hearts. Nuclear-encoded and mtDNA-encoded respiratory chain mRNA levels in heart tissues from control and Sod2-mKO mice were estimated through typical gene-level expression analysis of RNA-seq data. **A.** Heatmaps of nuclear-encoded subunit expression. The mRNAs are grouped by ETC complex and labeled with official gene symbols. Expression levels are reported as log_2_^FC^, where FC is fold-change relative to the control-group average. Green squares indicate higher expression levels, while red squares represents lower expression levels in SOD2-deficient hearts compared to the controls. Each column above “Individual mice” represents a single mouse; those above “Groups” represent the average values for three mice in each group. Differentially expressed genes are shown; the complete set of subunits with p values is provided in Supplementary Figure 1. **B.** Heatmap showing transcript levels of mtDNA-encoded respiratory chain subunits based on reference mitochondrial genome annotation with 13 mRNAs. No significant difference was detected in any gene. Heatmap and sample descriptions are the same as those provided in panel (A). **C.** The 11 mitochondrial mRNA transcripts represented in custom mtDNA annotation are shown with corresponding proteins. The bicistronic mRNA *mt-Atp8–6* is translated into ATP8 and ATP6; *mt-Nd4l–4* into ND4L and ND4. The other nine mRNAs encode one protein each. Complex I mRNAs and proteins are shown in shades of blue; Complex III in dark yellow, Complex IV in magenta; and Complex V (ATP synthase) in green. **D–F.** Total (D), complex-specific (E), and individual (F) levels of mtDNA-encoded mRNAs, based on the 11-mRNA custom mtDNA annotation (see Methods). Horizontal lines (D–E) and bars (F) show average values; circles (D–F) represent data from individual mice. The values are reported as multiples of the control mean (fold-control).

### 3.2. Annotation-agnostic analysis of RNA-seq data reveals increased abundance of mitochondrial tRNA sequences

We reasoned that gene-level expression analysis of SOD2-deficient hearts might have overlooked potential differences in the abundance of short intragenic and intergenic sequences in mitochondrial transcripts. Typical RNA-seq alignment procedures tolerate a small number of nucleotide mismatches. In addition, differential gene expression analyses estimate the expression level of each gene based on the *total* number of reads assigned to its annotated sequence. As a result, localized changes affecting short gene fragments may remain undetected, while potential changes outside annotated genes are completely excluded from gene-level quantification (Dobin et al., 2013; Engstrom et al., 2013). Such localized read-coverage information, which is lost through gene-level quantification, can be obtained using annotation-independent mapping approaches.

To investigate this possibility, we performed comprehensive 30-nt binning across the entire mtDNA and quantified each interval independently with the additional requirement of exact sequence matching (detailed in Methods). Consequently, we generated a mitochondrial genome-wide read-abundance profile at 30-nt resolution for a total of 544 short fragments, which provides more than 10x higher resolution compared to standard annotations with 37 sites (**fig 2A**). The control and SOD2-deficient groups exhibited largely overlapping profiles, corroborating the lack of significant differences found in gene-level mRNA expression. However, several regions showed clear divergence between the two profiles, indicating localized differences in read abundance (**fig 2A**, arrows).

Comparative analysis of these profiles identified a total of 17 short intervals with significantly altered abundance, which clustered into 10 distinct loci (**fig 2B**). Seven of these loci exhibited higher levels in SOD2-deficient samples and, without exception, coincided with tRNA genes. In contrast, none of the three loci showing lower abundance overlapped or neighbored a tRNA gene. Instead, two mapped to the 5’ termini of *mt-Cyb* and *mt-Co3* mRNA genes, which are among the three atypical mRNA transcripts that are not preceded by a tRNA gene (**fig 4C**). This latter observation may be relevant to the 5’-end maturation of mitochondrial mRNA transcripts lacking immediately upstream tRNAs but was not examined further in this study.

Overall, these results suggested an increased presence of mitochondrial tRNA sequences in SOD2-deficient heart RNA-seq data. High-throughput studies of tRNA expression require the use of specialized protocols for RNA isolation, pretreatment for removing nucleotide modifications, and customized library preparation to enable efficient detection of these small (about 70 nucleotides in length) RNA molecules (Zheng et al., 2015). Their capture through conventional RNA-seq protocols, which typically enrich for larger RNA molecules or specifically select polyadenylated transcripts, is not expected. Detection of mitochondrial tRNA sequences in conventional RNA-seq data has previously been reported in experimental models of defective RNA maturation and interpreted as evidence of tRNA–mRNA hybrid transcripts arising from incomplete RNA processing, rather than mature tRNAs (Jiang et al., 2019). In addition, mature tRNAs contain nucleotide modifications at several sites, making their detection unlikely with our approach requiring exact sequence matches. These suggested that tRNA sequences detected in SOD2-deficient hearts might be present as parts of long, hybrid molecules constituting immature polycistronic transcripts. We questioned whether these hybrid RNA species are biologically relevant and not exclusively found in engineered models with mitochondrial defects. Specifically, we asked whether intact polyadenylated RNA species containing tRNA sequences are present under physiological conditions in the healthy mouse heart and sought to characterize these hybrid species to facilitate the development of bioinformatic tools for their analysis.

**Figure 2.**
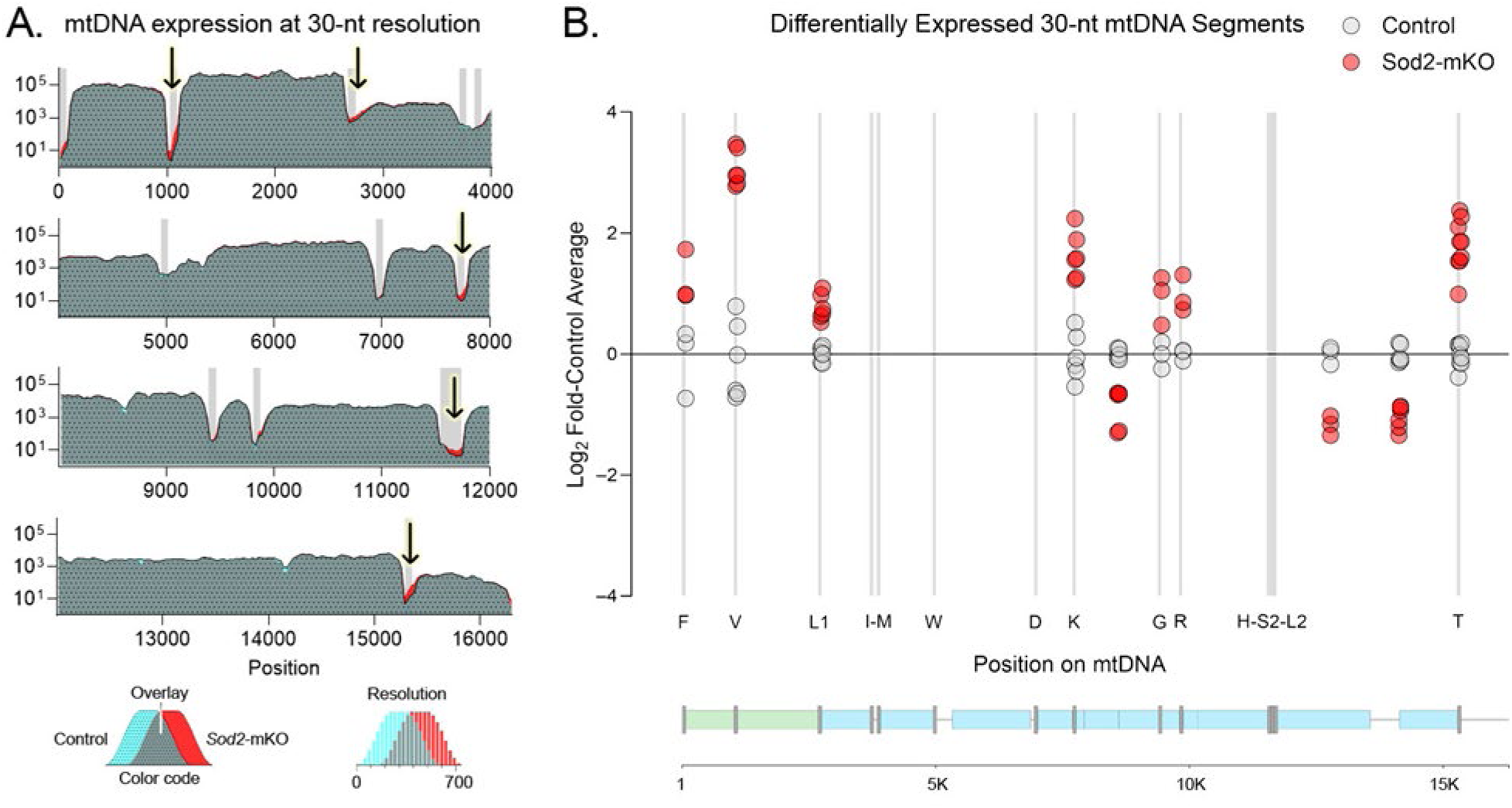
Annotation-agnostic analysis of mtDNA expression. Analysis of mitochondrial DNA heavy-strand expression at 30-nt resolution. **A.** The mouse mitochondrial genome (16,299 nt) is shown in three sections of 4K nt and a fourth section of 4,299 nt. A total of 544 fragments were quantified, and the mean values from three mice per group were plotted. The Y-axis shows abundance, expressed in counts per million mitochondrial reads, on a logarithmic scale. The x-axis indicates the genomic position of the midpoint of each 30-nt segment. The graph overlays the two groups, with light blue indicating controls; red, *Sod2*-mKO samples; and dark green, overlapping areas. Grey boxes mark tRNA sites. The arrows mark loci with an apparently higher abundance in *Sod2*-mKO samples. **B.** Differentially expressed 30-nt mtDNA fragments identified from the analysis of data summarized in panel (A). A total of 17 segments distributed among 10 loci were identified and plotted based on their mtDNA positions. Each symbol represents an individual mouse. Some positions appear to contain more than three data points per group because multiple adjacent segments were identified as differentially expressed. Y-axis shows log_2_^FC^, where FC is fold-change relative to the control mean. Vertical grey lines indicate tRNA gene positions, with the corresponding amino acids identified by single-letter codes along the x-axis. The x-axis is aligned with the mtDNA heavy-strand gene map shown below the graph. On the gene map, green boxes mark rRNA genes; blue, mRNA; and grey, tRNA. Sod2, superoxide dismutase 2; mKO, muscle-specific knockout; mtDNA, mitochondrial DNA; nt, nucleotide.

### 3.3. Direct sequencing of intact RNA reveals hybrid mitochondrial transcripts with polyA tails in healthy mouse heart

Direct long-read RNA sequencing is a relatively recent technology that has not replaced conventional short-read RNA-seq as the predominant approach for quantitative differential gene expression analysis, in part because of its lower throughput and more than tenfold higher error rate (Leshkowitz et al., 2022; Liu-Wei et al., 2024). However, by sequencing individual RNA molecules without prior fragmentation, direct RNA sequencing provides information on the sequence and structure of intact transcripts, including the physical linkage and organization of subregions within individual RNA molecules, making it particularly relevant for our purposes.

Previous studies reported accumulation of mtRNA maturation intermediates, including those containing tRNA fragments, primarily in experimental models with impaired RNA processing (Kuznetsova et al., 2017; Rackham et al., 2016; Siira et al., 2018). To determine whether tRNA sequences were present within intact polyadenylated RNA molecules under physiological conditions, we performed direct long-read RNA sequencing of healthy mouse hearts. First, we selected reads derived from mitochondrial transcripts and examined their length distributions. Compared to a typical RNA-seq experiment (150 bases, paired-end), which generated reads of length 150 bases or shorter, direct RNA sequencing generated reads spanning several thousand bases (**fig 3A**). The length distributions were nearly identical among the three mice. These included multiple peaks between 0.6 and 1.2 kilobases (kb) as well as prominent peaks at approximately 1.6, 2.4, and 3.5 kb (**fig 3A**), consistent with earlier analyses of polyadenylated mitochondrial RNA on northern blots (Bhat, Kantharaj, & Avadhani, 1984; Gelfand & Attardi, 1981).

Sequence analyses of individual mitochondrial transcripts revealed several types of hybrid species at substantial levels (**fig 3B**). Almost all tRNA sequences were present on polycistronic transcripts rather than existing as individual polyadenylated tRNA molecules. Transcripts containing mRNA sequences constituted about 53–56%. About one third of these also included tRNA fragments, which we characterized further (**fig 3C**). The majority consisted of a single mRNA gene accompanied by a single tRNA. We also detected, albeit much less frequently, even longer polycistronic RNA molecules containing multiple mRNA and tRNA fragments, some of which also included rRNAs. These findings confirm the presence of tRNA sequences in polyadenylated RNA species, corroborating recent findings (McShane et al., 2024), and reveal multiple forms of mRNA-containing transcripts retaining uncleaved tRNA segments in healthy mouse hearts.

Importantly, species containing mRNA sequences together with uncleaved tRNA extensions were not merely unprocessed transcripts but partially processed intermediates as they were cleaved and polyadenylated at their 3’ ends. The presence of the polyA tail is critical as it leads to their incorrect classification as mature mRNAs in short-read RNA-seq studies, hindering their identification as immature RNA species.

Although direct RNA sequencing enabled us to obtain an initial characterization of these immature mitochondrial RNA species, most publicly available RNA-seq datasets, including the Sod2-mKO dataset analyzed in this study, have been generated using conventional short-read sequencing. To enable analysis of these incompletely processed RNA species in existing short-read datasets, we set out to develop a new bioinformatics tool.

**Figure 3.**
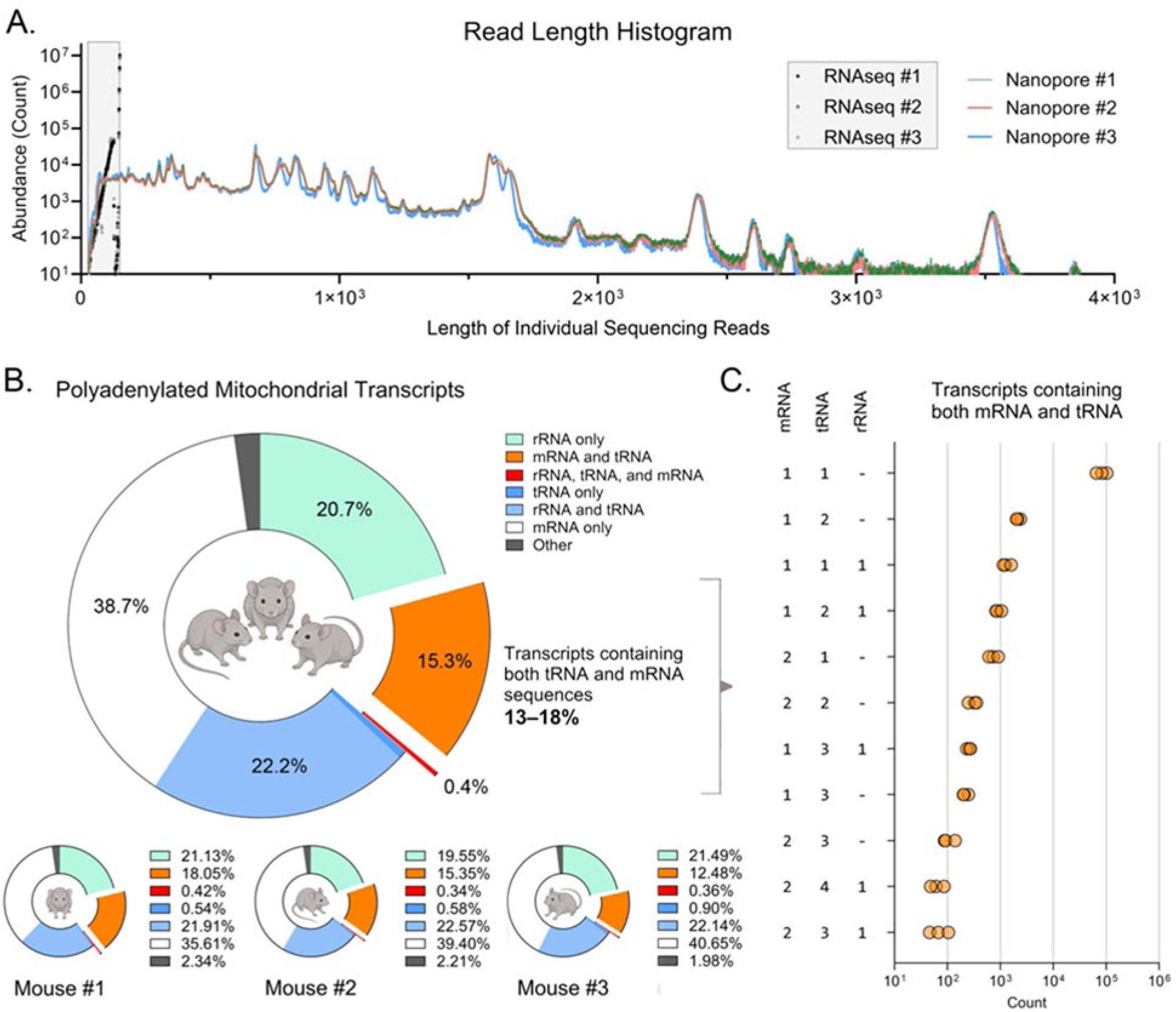
Polyadenylated but immature RNAs in healthy mouse heart examined through direct RNA sequencing. **A.** Histogram of sequencing read lengths from direct RNA sequencing (Oxford Nanopore) of wild-type mice compared to those from short-read RNA-seq (Illumina) of control mice in the Sod2-mKO dataset described earlier. The y-axes shows absolute numbers of reads with each length on a logarithmic scale. Dots colored in shades of grey mark short-read RNA-seq reads obtained from three control mice. The grey box marks the area covered by these reads, with read lengths ranging from 30 nt to 150 nt. Green, blue, and red dots indicate direct RNA sequencing reads from three wild-type mice, with read lengths of several thousand nucleotides. Note the consistent distribution pattern among the three mice, with peaks appearing at sizes described in earlier studies of polyadenylated mtRNAs. **B.** Distribution of polyadenylated mitochondrial RNAs classified by content. Direct long-read RNA sequencing data were utilized to analyze intact polyadenylated mitochondrial transcripts. Estimated ratios of mitochondrial polyadenylated RNA types are shown on a pie chart, with average values from three mice. Values from individual mice are shown below in smaller pie charts. **C.** Abundance of subsets of polyadenylated mitochondrial RNAs containing both messenger and transfer RNAs based on direct RNA sequencing. The subsets are defined by the numbers of distinct mRNA, tRNA, and rRNA species found in a single read and shown on the y-axis. The x-axis shows the number of reads in each read set on a logarithmic scale. Circles represent data from individual mice (n = 3).

### 3.4. Development of a new tool for quantitative imRNA analysis using short-read RNA-seq datasets

Nuclear mRNA maturation is highly coordinated, with 5’ capping occurring early during pre-mRNA processing, followed by 3’-end cleavage and polyadenylation (Moore & Proudfoot, 2009). Consequently, polyadenylated transcripts retaining unremoved extensions are not expected nuclear RNA species. Unlike nuclear genes, mitochondrial genes are not transcribed as individual units but as long polycistronic RNAs containing multiple rRNA, tRNA, and mRNA sequences, which are then separated and processed, yielding mature RNA species (Aloni & Attardi, 1971; Anderson et al., 1981; Mercer et al., 2011; Ojala, Montoya, & Attardi, 1981). Importantly, mitochondrial RNA processing does not necessarily proceed in a strict order among junctions: processing and polyadenylation of the 3’ end of an RNA molecule can occur while its upstream sequences remain uncleaved, generating a partially processed RNA containing uncleaved extensions (Kuznetsova et al., 2017; McShane et al., 2024; Mercer et al., 2011) (**fig 4A, fig 3B–C**).

#### Identification of immature mRNAs

We define an **immature mRNA (imRNA)** of a protein-coding gene as an mRNA-like molecule that contains (i) the complete coding sequence of its cognate mature mRNA, together with (ii) an unremoved tRNA sequence extension retained due to incomplete processing, and (iii) a 3’ polyA tail (**fig 4B**). Thus, although an imRNA is polyadenylated and contains the complete coding sequence present in a mature mRNA, it retains uncleaved sequences that distinguish it from the mature transcript. Unlike mature mRNAs, imRNAs are substrates for RNA processing rather than translation.

Conventional gene-level RNA-seq strategies have been developed based on nuclear RNA biology, with the underlying assumption that polyA-selected RNA pools are highly enriched in mature mRNAs. The quantification algorithms rely on short read segments that match mRNA sequences and cannot distinguish imRNAs from their cognate mature mRNAs. Conceivably, reads spanning an uncleaved gene boundary can provide sequence information specific to incompletely processed molecules and therefore could be utilized to extract information on imRNA from short-read RNA-seq datasets through computational analyses. However, existing pipelines for high-throughput sequencing analysis do not quantify reads spanning uncleaved mitochondrial gene boundaries, leaving data in short-read datasets unexamined. We therefore developed an *in silico* analysis tool for imRNA analysis that identifies reads spanning mitochondrial tRNA–mRNA and mRNA–tRNA boundaries to estimate imRNA levels.

#### Uncleaved junction identifiers

We focused on two types of uncleaved junctions that could serve as indicators of imRNA molecules: (i) tRNA–mRNA junctions, indicating an imRNA retaining the upstream tRNA segment (hereafter, imRNA with 5’ tRNA); and (ii) mRNA–tRNA junctions, indicating an imRNA retaining the uncleaved downstream tRNA (hereafter, imRNA with 3’ tRNA). This distinction reflects the underlying mechanism of mtRNA maturation: tRNA–mRNA junctions are cleaved by an endonuclease, RNase Z, that processes 3’ ends of tRNAs, while mRNA–tRNA junctions are cleaved by a multiprotein complex, RNase P, that processes 5’ ends of tRNAs.

Based on the current gene annotation, we identified seven tRNA–mRNA junctions and seven mRNA–tRNA junctions within the polycistronic transcript originating from the heavy-strand promoter (**fig 4C**). For each junction, we generated uncleaved junction identifier (UJI; detailed in Methods) candidates and subjected them to a validation procedure as described below to ensure specific detection of intended mitochondrial transcripts.

#### Specific detection of mtDNA transcripts despite nuclear-encoded mitochondrial DNA segments (NUMTs)

A major shortcoming of short-read sequencing is the difficulty of confidently assigning reads to their loci of origin when multiple loci share highly similar sequences (Treangen & Salzberg, 2011), which is relevant for mitochondrial transcripts (Maude et al., 2019). Nuclear genomes contain numerous sequences with high similarity to mtDNA, known as nuclear-encoded mtDNA segments (NUMTs), which are thought to originate from the integration of mtDNA fragments into nuclear chromosomes (Bar-Yaacov et al., 2013; Richly & Leister, 2004; Wei et al., 2022). NUMTs complicate the confident assignment of certain mtDNA-encoded sequences to mitochondrial transcripts. We address this complication through computational and biological validation, to demonstrate that any contribution to detected UJIs from nuclear transcripts is minimal to nonexistent.

Through *in silico* analyses, we identified 44 NUMTs distributed across 15 mouse chromosomes (**fig 4D**). For seven of the 14 mouse gene junctions, we were able to assign specific UJIs that were not present in NUMTs. For the remaining junctions, all potential UJI sequences were also present in the nuclear genome in three mouse NUMT regions: NUMT-1 (NC_000067.7, chr1:24650616–24655269), NUMT-2 (NC_000068.8, chr2:22477300–22480546), and NUMT-3 (NC_000068.8, chr2:69185882–69186010). We therefore examined whether transcription from these loci could confound the specific detection of mtDNA-encoded transcripts. Inspection of genomic RNA-seq coverage across these regions in mouse heart, kidney, liver, skeletal muscle, spleen, and brain revealed abnormal coverage boundaries precisely coinciding with the 5’ and 3’ boundaries of the mtDNA-matching regions (**fig 4F**). This profile is distinct from the tapered coverage observed at the boundaries of biological transcripts, suggesting artificial mapping of mtRNA sequences to NUMTs rather than genuine transcription from these loci. We therefore selected the remaining seven UJIs from these regions, resulting in a set of 14 mouse UJIs for subsequent biological validation (Supplementary Table 1).

In humans, we identified 188 NUMTs distributed across 24 chromosomes and three unplaced scaffolds (**fig 4E**). For eight of the 14 human junctions, we were able to select mtDNA-specific UJIs that were absent from human NUMTs. For the remaining six junctions, candidate UJIs were also present in three nuclear regions: NUMT-1 (NC_000001.11, chr1, 629639–629668), NUMT-2 (NC_000002.12, chr2, 87824941–87824970), and NUMT-3 (NC_000005.10, chr5, 60761585–60761556). Assessment of potential transcription of these three regions using the same RNA-seq coverage analysis described above for mice revealed similar coverage patterns, consistent with mitochondrial reads mapping to homologous nuclear sequences. Accordingly, we included the six UJIs present in these regions and obtained a set of 14 human UJIs with presumed specificity for subsequent biological validation (Supplementary Table 2).

In summary, our matching algorithm required each UJI to match a continuous 30-nt sequence with 100% identity, disqualifying partial, interrupted, or shorter nuclear matches. This stringent criterion ruled out most NUMT-derived sequences as potential sources of false-positive matches and, together with the RNA-seq coverage analysis described above, led to the selection of 14 UJIs per species predicted to specifically detect mitochondrial uncleaved junctions pending their biological validation.

**Figure 4.**
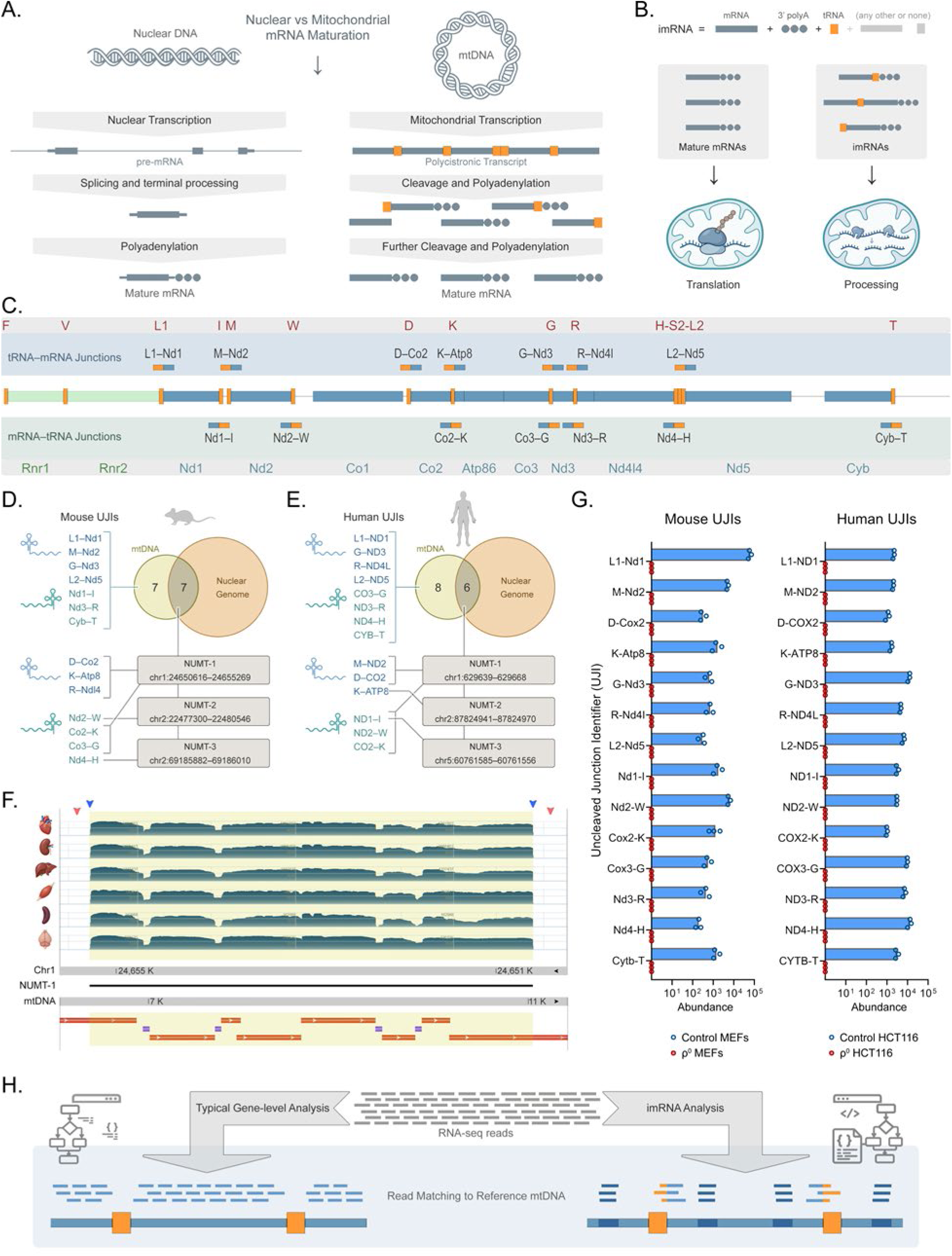
Development of a new tool for quantitative imRNA analysis using short-read RNA-seq datasets. **A.** Comparison of nuclear and mitochondrial mRNA maturation and their processing intermediates. **B.** Illustration of the definition of imRNA and its functional distinction from mature mRNAs. **C.** Positions of uncleaved junction identifiers (UJIs). Mouse mitochondrial DNA is shown in the center. Blue segments indicate protein-coding regions; orange segments, tRNAs; light green segments, rRNAs. Transfer RNA genes are labeled on the upper grey band using single-letter codes for their cognate amino acids (dark red). Ribosomal RNA (green) and mRNA (blue) genes are labeled on the lower grey band. For simplicity, the “mt-” prefixes are omitted from gene symbols. The blue band above the genome map marks the seven UJIs spanning tRNA–mRNA junctions; the green band below the map marks the seven UJIs spanning mRNA-tRNA junctions. Each UJI contains a tRNA-matching segment (orange) and an mRNA-matching segment (blue). UJIs are not drawn to scale to improve visibility. **D.** Presence of candidate mouse UJIs in the mitochondrial and nuclear genomes. Of the 14 UJIs, seven were found exclusively in the mitochondrial genome; the remaining seven were also present at three nuclear loci on chromosomes 1 and 2. **E.** Presence of candidate human UJIs in the mitochondrial and nuclear genomes. Of the 14 UJIs, eight were found exclusively in the mitochondrial genome; the remaining six were also present at three nuclear loci on chromosomes 1, 2, and 5. The UJI *ND1–I* is present at two nuclear loci, one on chromosome 1 and the other on chromosome 5. **F.** Assessment of UJI-containing NUMT expression in mouse tissues. Mouse NUMT-1 coverage map is shown as a representative example of the three mouse NUMTs containing UJIs. NUMT-1 (horizontal black line; chr1:24,650,616– 24,655,269, negative strand; mtDNA:6390–11042) is shown aligned to mouse chromosome 1 (above; 24,650,316– 24,655,569, negative strand) and mtDNA (below; 5390–11042). The latter two include 300-nt flanking sequences on both sides. Numbers on grey bands mark genomic positions. Yellow shading marks the NUMT-aligned regions; white regions indicate flanking sites. The six tracks on the nuclear genome show RNA-seq read coverage in heart, kidney, liver, skeletal muscle, spleen, and brain, from top to bottom. Mitochondrial mRNA (red bars) and tRNA (purple bars) sites are indicated. Note the sharp drops in read coverage at both ends of NUMT-1 (blue arrows), with no reads mapping to the flanking sites that differ in sequence from mtDNA (red arrows). Coverage also decreases over tRNA sequences, consistent with their removal during mitochondrial RNA processing, suggesting that the mapped reads were derived from mitochondrial transcripts. Together, these patterns support mismapping of mitochondrial transcripts to NUMT-1 rather than transcription of this nuclear locus. **G.** Specificity analyses of mouse and human UJIs using cells lacking mitochondrial transcripts. Mouse RNA-seq data were obtained from mouse embryonic fibroblast (MEF) cultures with intact mtDNA (Control MEFs; n = 3) and ρ^0^ MEFs (n = 3). Human RNA-seq data was obtained from human colon tumor cell (HCT116) cultures with intact mtDNA (Control HCT116; n = 3) and ρ^0^ HCT116 cells (n = 3). In both graphs, UJI levels are shown on the x-axis as counts per billion reads. The 14 UJIs are labeled using the pair of junction-flanking genes in 5’-to-3’ order. For brevity, mitochondrial tRNA genes are represented using the standard single-letter codes of their cognate amino acids rather than official gene symbols. A complete list of UJIs, with gene names and official gene symbols, is provided in Supplementary Tables 1–2. The bars indicate mean values; dots represent biological replicates. Counts in all ρ^0^ samples (red symbols) were zero; no UJI was detected in either mouse or human cells lacking mitochondrial mRNA. ρ^0^ cells, cells lacking mtDNA. **H.** Strategy for imRNA analysis using short-read RNA-seq datasets, compared to gene-level analysis. In conventional gene-level analysis (left), reads assigned to annotated exons are counted to estimate gene expression levels. In imRNA analysis (right), sequences matching predefined junction-spanning sites and reference mRNA segments (dark blue) are identified and counted to estimate relative imRNA abundance.

#### Biological validation of specificity of selected UJIs for imRNA quantification

Computational analyses based on reference sequences as described above might not account for imperfections present in real-world RNA-seq data, such as genomic variants and sequencing errors. In addition, nuclear RNA processing, including splicing and nucleotide editing, could conceivably generate transcripts containing UJIs that are not present in the genomic DNA. It is therefore critical to determine the relative abundance of such potential nuclear transcripts compared with UJIs authentically present in mitochondrial transcripts.

Mammalian cells can be cultured under certain conditions to eliminate mtDNA, resulting in the generation of ρ^0^ (rho-zero) cells devoid of mitochondrial DNA and consequently unable to produce mitochondrial RNA (Desjardins, Frost, & Morais, 1985; Martinus et al., 1996). We reasoned that detection of UJIs in ρ^0^ cells lacking mitochondrial transcripts would indicate misidentification of NUMT-derived RNA molecules as mitochondrial RNA. We utilized publicly available data from an RNA-seq study (SRP134850, see Methods; supplementary table 6) that included both control and ρ^0^ MEFs (Shimada et al., 2018) to test the specificity of the 14 mouse UJIs for mitochondrial transcripts. The UJIs were consistently detected in the control cells but not in ρ^0^ cells, with zero matches in the mtDNA-deficient group, suggesting that these UJIs specifically detect mitochondrial transcripts despite the presence of NUMTs in the mouse genome (**fig 4G**, left; supplementary table 3).

We performed a similar analysis for the human UJIs using an RNA-seq dataset (SRP093357, see Methods; supplementary table 6) containing both control and mtDNA-deficient human colon tumor cells (Heller et al., 2017). Cancer cell lines, in contrast to primary cells, typically accumulate mutations, increasing the chances of emergence of mtDNA-matching short fragments in the nuclear genome. However, no reads containing any of the 14 human UJIs were found in ρ^0^ cells, while all were detected in the controls (**fig 4G**, right; supplementary table 4). These data confirm that the 14 human UJIs may be used for the specific detection of mitochondrial RNAs while excluding nuclear transcripts.

Furthermore, the UJI counts obtained in control cells suggest that (i) imRNAs are present in mouse and human cells cultured in standard conditions without a particular treatment or manipulation, and (ii) existing RNA-seq datasets generated using earlier sequencing technologies and varying read layouts, read lengths, and sequencing depths (see Methods) can be utilized to detect UJIs to estimate imRNA levels.

#### Quantification of imRNAs for comparison between samples with varying abundance of mitochondrial RNA

Comparison of RNA-seq-derived measures between samples requires normalization for both biological and technical variation within individual experiments. The common practice of expressing gene-specific read counts as a portion of total read counts in RNA-seq analysis is not ideal for imRNA quantification for several reasons. For instance, an overall increase in mitochondrial abundance and hence transcription could result in an apparent increase in imRNA levels expressed as a fraction of total transcriptome reads, while a decrease in total mtRNA abundance could generate the opposite effect. Normalizing to total mitochondrial read counts, on the other hand, could be misleading under conditions that induce nonuniform effects on individual transcripts. Therefore, we concluded that normalization of each imRNA species to its cognate mRNA provided the most robust and biologically relevant measure of relative imRNA abundance (**fig 4H**).

### 3.5. Levels of imRNAs with 5’ tRNA are elevated in SOD2-deficient hearts

We estimated the levels of the 14 imRNA species in control and SOD2-deficient mouse hearts. Total imRNA levels did not differ between the two groups (**fig 5A**). However, separate analysis of the two types suggested a specific increase in imRNAs retaining 5’ tRNA (**fig 5B**). Analysis of individual imRNA species further revealed significantly elevated levels of imRNAs with uncleaved tRNA–mRNA junctions (imRNAs with 5’ tRNA) in SOD2-deficient hearts (**fig 5C**). In contrast, no increases were detected in imRNAs with mRNA-tRNA junctions (imRNAs with 3’ tRNA) (**fig 5D**). Increased mtROS and oxidative stress have been well established in SOD2-deficient models, and were confirmed in the SOD2-mKO cohort used in the generation of RNA-seq data analyzed here (Kauppila et al., 2018). In addition, we found that the overall mitochondrial transcriptome or individual mtDNA-encoded genes show no decreases in typical gene-level RNA-seq analysis (**fig 1B–F**). Taken together, the observed pattern of imRNA levels in control and SOD2-deficient hearts suggest that mtROS might induce a specific effect in the form of elevated imRNAs with 5’ tRNA as opposed to a general defect in RNA maturation or a broader deficiency in mitochondrial RNA metabolism.

### 3.6. Elevation of imRNAs with 5’ tRNA extensions as a potential indicator of high mtROS

This distinctive imRNA signature, characterized by increased levels of imRNAs retaining 5’ tRNAs but not those containing 3’ tRNAs, suggested that mtROS may drive this phenotype. However, it is plausible that other types of mitochondrial perturbation could trigger similar effects. We therefore asked whether this imRNA pattern could be induced by other forms of mitochondrial stress or represent a specific signature of increased mtROS.

#### *Mterf4*-mKO mice with mitochondrial translation deficiency and mtRNA accumulation

We reasoned that our analysis could generate false positives in samples with substantially elevated mRNA abundance, which could overwhelm nascent RNA processing and increase imRNA levels. Accordingly, we sought a model of mitochondrial stress that features both mitochondrial dysfunction and mtRNA accumulation. Mitochondrial transcription termination factor 4 (*Mterf4*), although its original name suggests transcriptional regulation, is required for mitochondrial ribosome assembly and translation. While global *Mterf4* knockout is lethal, mice with muscle-specific *Mterf4* deletion (*Mterf4*-mKO) survive up to 4–5 months and develop mitochondrial cardiomyopathy. Notably, these mice also show elevated mtDNA and mtRNA levels, making them a model of severe mitochondrial stress accompanied by mtRNA accumulation.

Our analysis of RNA-seq data from *Mterf4*-mKO mice confirmed an overall increase in mitochondrial transcript levels in MTERF4-deficient hearts (**fig 6A**, dot plot), with a more than twofold increase in multiple mtRNA species, as reported previously (Cámara et al., 2011). Despite this accumulation, we found no increase in imRNA levels (**fig 6A**, bar graphs). Instead, we detected significant decreases in certain imRNA species of both types (M–Nd2, Nd1–I, and Nd2–W). The absence of imRNA elevation suggests that robust increases in total mitochondrial DNA and RNA levels, despite causing severe mitochondrial dysfunction, do not induce the specific imRNA pattern observed in SOD2 deficiency.

#### *Tfam*-mKO and *Polrmt*-mKO mice with drastic mtRNA deficiency

More severe forms of mitochondrial stress, characterized by complete loss of aerobic respiration, can be induced by suppressing mitochondrial RNA synthesis. Mitochondrial transcription factor A (TFAM) and mitochondrial RNA polymerase (POLRMT) are both required for mitochondrial transcription. In mice, global knockout of either gene causes embryonic death (Kühl et al., 2016; Larsson et al., 1998). Muscle-specific knockout (mKO) models are viable, although they have short lifespans. Both models exhibit decreased mtRNA levels, leading to reduced ETC function and mitochondrial cardiomyopathy. We analyzed RNA-seq datasets from heart tissue of *Tfam*-mKO and *Polrmt*-mKO mice (Kühl et al., 2017) to assess the mitochondrial transcriptome and imRNAs. We confirmed a substantial decline in mtRNA in both models. *Tfam* deletion reduced mitochondrial mRNA abundance to below 10% of control levels (**fig 6B**, dot plot). *Polrmt* deletion caused a more pronounced defect, with an almost complete loss of the mitochondrial transcriptome (**fig 6C**, dot plot). In *Tfam*-mKO samples, levels of imRNAs containing 3’ tRNAs were also lower, with multiple species below the detection limit. Levels of imRNAs containing 5’ tRNA did not mirror this decline, nor did they show a significant increase (**fig 6B**, bar graphs). Of note, one of the 14 imRNA species showed an apparent increase: mean *K–Atp8* imRNA levels were about twice those in controls (p = 0.052). In *Polrmt*-mKO samples, nearly all imRNAs exhibited lower levels, with the exception of the imRNA *L1–Nd1*, which was not altered (**fig 6C**, bar graphs). These findings suggest that severe mitochondrial dysfunction alone does not induce the pattern of imRNA elevation observed in the model with elevated mtROS.

### *Lonp1*-smKO mice with mitochondrial proteotoxic stress

A mouse model of mitochondrial proteotoxic stress, generated by muscle-specific deletion of *Lonp1* encoding the mitochondrial protease LONP1 (lon peptidase 1, mitochondrial), showed no indications of a change in the levels of imRNAs retaining 5’ tRNAs or those with 3’ tRNAs (**fig 6D**). This dataset consisted of two mice per group; additional data are needed to investigate imRNA status in LONP1-deficient tissues.

### *Ant1*-overexpressing mice as a model of mitochondrial stress with reduced mtROS

Lastly, we sought a mouse model of perturbed mitochondrial homeostasis accompanied by a demonstrated lack of mtROS elevation. We specifically searched for studies that performed both ROS assessments and RNA sequencing on the same experimental model. Adenine nucleotide translocase 1 (ANT1) is a nuclear-encoded protein located in the mitochondrial inner membrane where it mediates ATP/ADP exchange. ANT1 overexpression can clog mitochondrial import machinery and disrupt the mitochondrial proteome, resulting in a progressive reduction in respiration (X. Wang et al., 2022). Transgenic mice overexpressing *Ant1* (*Ant1*^Tg/+^ mice), a model of mitochondrial import stress, exhibit muscle atrophy and reduced lean mass, but no evidence of ROS elevation; instead, mitochondria isolated from *Ant1*^Tg/+^ mice show reduced H_2_O_2_ production (Brennan et al., 2026). This model therefore was deemed suitable to test the specificity of the imRNA pattern as an indicator of mtROS stress (**fig 7A**).

We analyzed the RNA-seq data generated from skeletal muscle of *Ant1*^Tg/+^ mice (Brennan et al., 2026). ANT1 overexpression did not cause a significant change in mitochondrial mRNA levels (**fig 7B**). Neither class of imRNAs showed increased abundance. Instead, the total abundance of imRNAs retaining 5’ tRNA **(fig 7C)** and the levels of individual imRNAs *M–Nd2* and *D–Co2* (**fig 7D**) were significantly reduced, opposite to the increases observed in SOD2-deficient hearts. This reduction in imRNAs with 5’ tRNA is consistent with a potential association between mtROS levels and the abundance of these processing intermediates.

Overall, comparison across six mouse models exhibiting several forms of mitochondrial stress suggests that increased abundance of imRNAs retaining 5’ tRNA might be a specific indicator of elevated mitochondrial ROS.

### 3.7. Rotenone-treated human iPSCs show an imRNA pattern consistent with a conserved response

We asked whether the mtROS-induced imRNA elevation observed in SOD2-deficient mouse hearts was also detectible in human cells with elevated mtROS. Cells treated with rotenone, an inhibitor of the ETC Complex I, provide a pharmacological model of mitochondrial ROS elevation (**fig 8A**). We analyzed RNA-seq data (NCBI BioProject, PRJNA1188018) from human induced pluripotent stem cells derived from umbilical cord cells (Fujioka, Shimizu, Yoshino, Miyoshi, & Nakamura, 2010) and treated with increasing doses of rotenone (Hongen et al., 2024). Gene-level analysis suggested potential increases in *ND1* and *CO1* mRNA levels but no changes in the other mitochondrial mRNAs or in total mitochondrial mRNA abundance (**fig 8B**). In contrast, imRNA analysis revealed an overall increase in imRNAs with 5’ tRNA, with no evidence of increased abundance of imRNAs with 3’ tRNA (**fig 8C–E**). These results from human iPSCs largely reproduced the pattern observed in SOD2-deficient mouse hearts (**fig 5**), characterized by a specific elevation of imRNAs retaining 5’ tRNA extensions.

**Figure 5.**
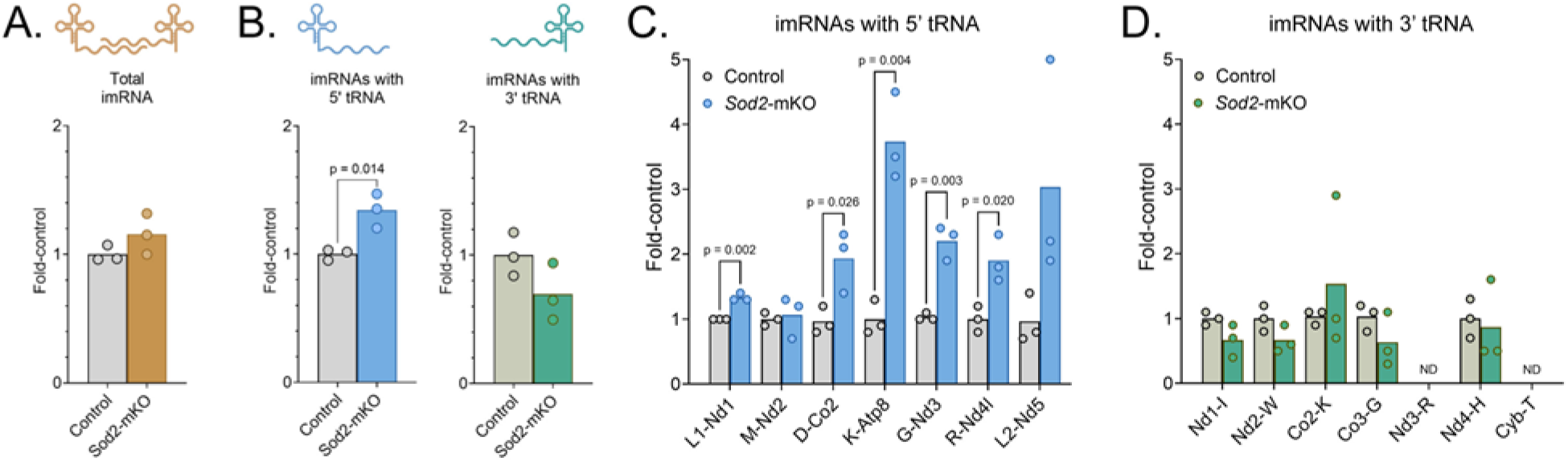
Analysis of imRNAs in SOD2-deficient hearts Mitochondrial imRNA analysis of hearts from *Sod2*-mKO (n = 3) and control (n = 3) mice. **A.** Total imRNA abundance. **B.** Combined levels of imRNAs with 5’ tRNAs, and those with 3’ tRNAs. **C.** Levels of individual imRNAs with 5’ tRNA (mKO group in blue). **D.** Levels of individual imRNAs with 3’ tRNA (mKO group in green). In all panels, imRNA levels were estimated using the imRNA analysis tool developed in this study (see Methods) and expressed as multiples of the control mean (fold-control). For brevity, mitochondrial tRNA genes were denoted by single-letter symbols of corresponding amino acids; mRNA genes by shortened gene symbols without the “mt-” prefix. Individual imRNAs are indicated on the x-axes with the uncleaved junction identifiers (UJIs) they contain, *e.g.*, L1–Nd1 marks the *imRNA mt-Nd1 retaining a 5’ mt-Tl1 extension*. Mouse UJI descriptions with official gene names are provided in Supplementary Table 1. Bars show mean values; circles represent individual mice. Significant differences are marked on the graphs with corrected p-values for multiple comparisons. imRNA, immature mRNA; mKO, muscle-specific knockout; ND, not detected.

## 4. Discussion

### Two main contributions

Our study provides two complementary contributions to the investigation of mitochondrial RNA metabolism. First, we developed a computational tool for estimating mitochondrial immature mRNA (imRNA) levels using conventional short-read RNA-seq data. This approach distinguishes changes in uncleaved mitochondrial gene junctions that are obscured by gene-level expression estimates and enables retrospective analysis of existing datasets. Second, using this tool, we identified an imRNA pattern potentially associated with mitochondrial oxidative stress: elevated levels of imRNAs retaining 5’ tRNA extensions without corresponding increases in those retaining 3’ tRNA extensions. This pattern was observed in SOD2-deficient mouse hearts (**fig 5**) and largely reproduced in rotenone-treated human cells (**fig 8**) but not in other mitochondrial stress models (**figs 6–7**). Our work strongly suggests that mitochondrial RNA maturation intermediates contain information about mitochondrial dyshomeostasis that is not captured by conventional transcriptome analyses but can be extracted from existing RNA-seq datasets through imRNA analysis.

**Figure 6.**
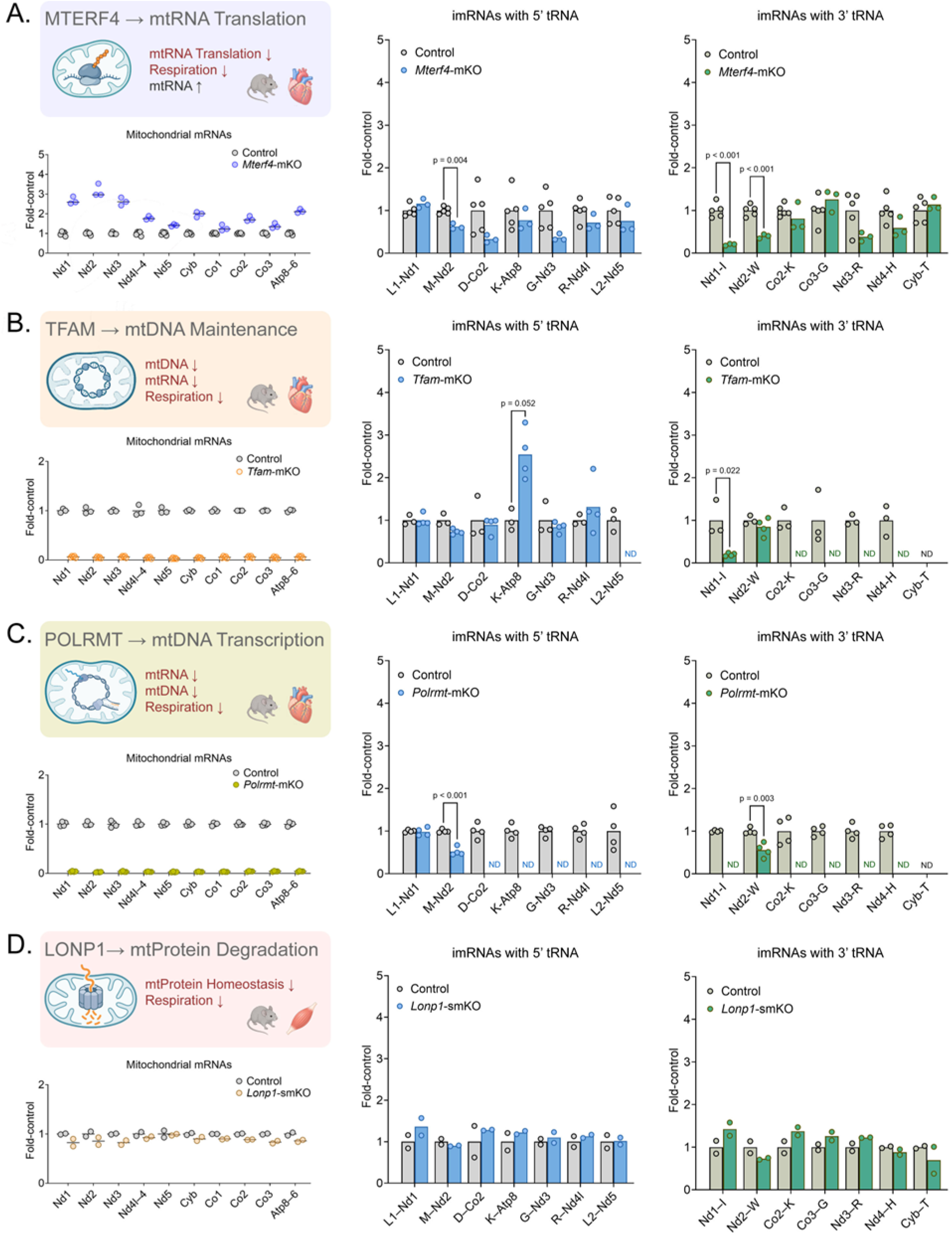
Mitochondrial mRNA expression and imRNA analysis in mouse models of mitochondrial stress Mitochondrial mRNA and imRNA analyses of tissue samples from mice with mitochondrial deficiencies. **A.** *Mterf4*-mKO (n = 3) and control (n = 5) mice. **B.** *Tfam*-mKO (n = 4) and control (n = 3) mice. **C.** *Polrmt*-mKO (n = 4) and control (n = 4) mice. **D.** *Lonp1*-mKO (n = 2) and control (n = 2) mice. Mitochondrial mRNA levels are shown on dot graphs. *Nd4l–4*, bicistronic transcript encoding both *mt-Nd4l* and *mt-Nd4*; *Atp8–6*, bicistronic transcript containing both *mt-Atp8* and *mt-Atp6*. Results of imRNA analyses are presented on pairs of bar graphs as indicated. Relative levels of imRNAs with 5’ tRNA (left, mKO group in blue) and those with 3’ tRNA (right, mKO group in green) were estimated using the imRNA analysis tool developed in this study (see Methods). Individual imRNAs are indicated on the x-axes with the uncleaved junction identifiers (UJIs) they contain. For brevity, mitochondrial tRNA genes were denoted by single-letter symbols of corresponding amino acids; mRNA genes by shortened gene symbols without the “*mt*-” prefix. Mouse UJI descriptions and official gene names are provided in Supplementary Table 1. Y-axis limits were set to five to facilitate comparison to Sod2-mKO results (fig 5). In all panels, the values were normalized to the average of controls (grey bars) and expressed as multiples of the control mean (fold-control). Lines and bars show mean values; circles represent individual mice. Significant differences are marked on the graphs with corrected p-values for multiple comparisons; p values below 0.001 were indicated as p < 0.001 rather than as exact values. Statistical analysis was not performed for *Lonp1*-smKO data as only two biological replicates were available. imRNA, immature mRNA; mKO, muscle-specific knockout; smKO, skeletal muscle-specific knockout; UJI, uncleaved junction identifier.

**Figure 7.**
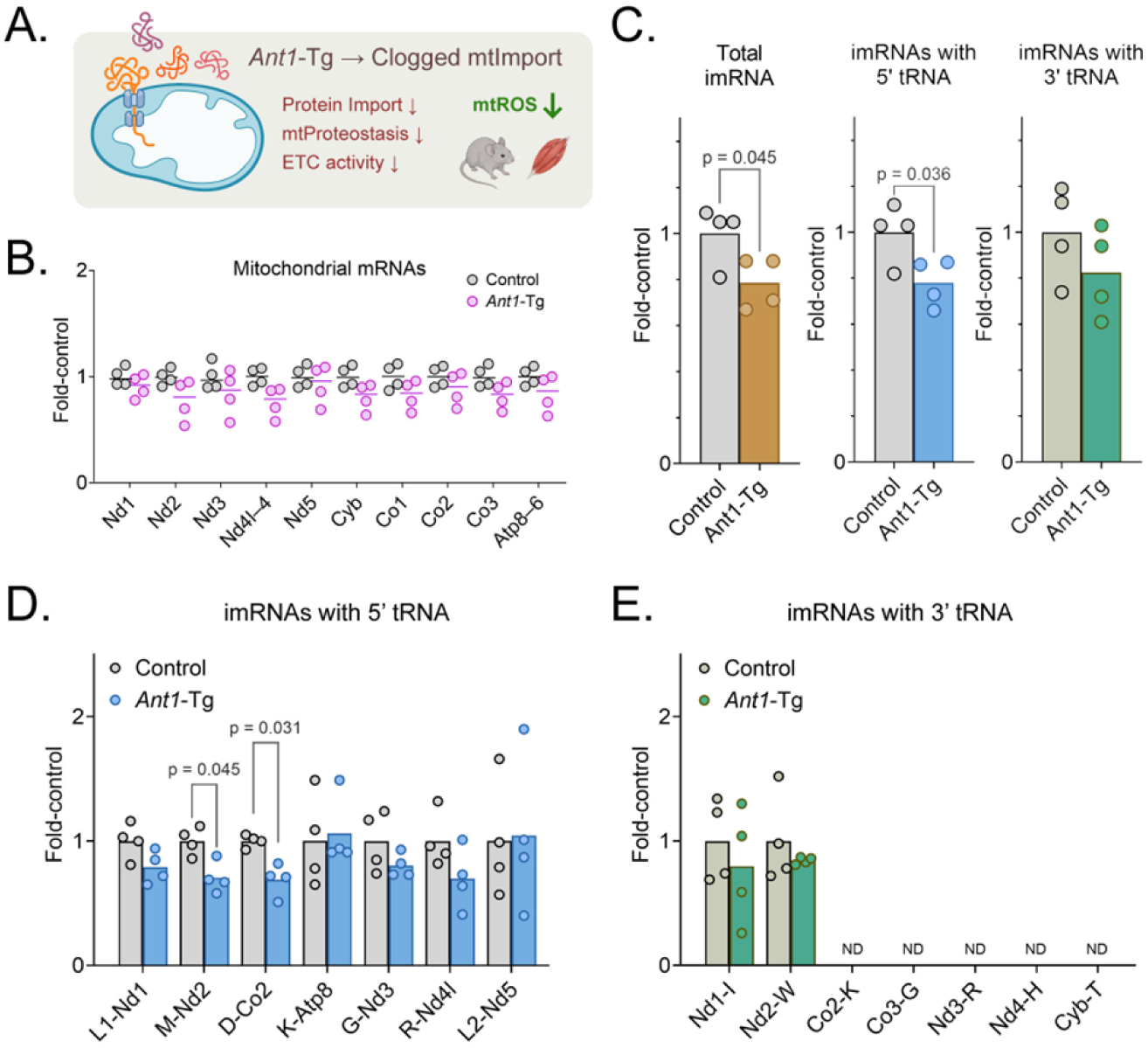
Mitochondrial mRNA levels and imRNA analysis in a mitochondrial stress model with reduced mtROS. Mitochondrial mRNA and imRNA analyses of quadriceps skeletal muscle samples from transgenic mice overexpressing *Ant1* (*Ant1*-Tg; n = 4) and wild-type mice (Control; n = 4). **A.** ANT1 overexpression clogs the mitochondrial protein import machinery, disrupts proteome homeostasis, decreases respiration, and reduces mitochondrial ROS production. **B.** Mitochondrial mRNA levels are shown on a dot plot. *Nd4l–4*, bicistronic transcript encoding *mt-Nd4l* and *mt-Nd4*; *Atp8–6*, bicistronic transcript containing *mt-Atp8* and *mt-Atp6*. Genes are labeled on the x-axis using shortened gene symbols without the “*mt*-” prefix. **C.** Total imRNA abundance and cumulative abundance of imRNAs retaining 5’ or 3’ tRNAs. **D–E.** Estimated levels of individual imRNAs with 5’ tRNA (D) and with 3’ tRNA (E). Individual imRNAs are labeled on the x-axes with the uncleaved junction identifiers (UJIs) they contain; for instance, L1–Nd1 denotes an Nd1 imRNA retaining a 5’ tRNA L1 extension. For brevity, mitochondrial tRNA genes are denoted by the single-letter symbols of corresponding amino acids, and mRNA genes by shortened symbols without the “*mt*-” prefix. Mouse UJI descriptions and official gene names are provided in Supplementary Table 1. In all graphs, values were normalized to the control mean and expressed as multiples of the control mean (fold-control). Lines and bars show mean values; circles represent individual mice. Significant differences are marked on the graphs with p values corrected for multiple comparisons. Abbreviations: *Ant1*-Tg, *Ant1*-overexpressing transgenic mice; imRNA, immature mRNA; ND, not detected; UJI, uncleaved junction identifier.

**Figure 8.**
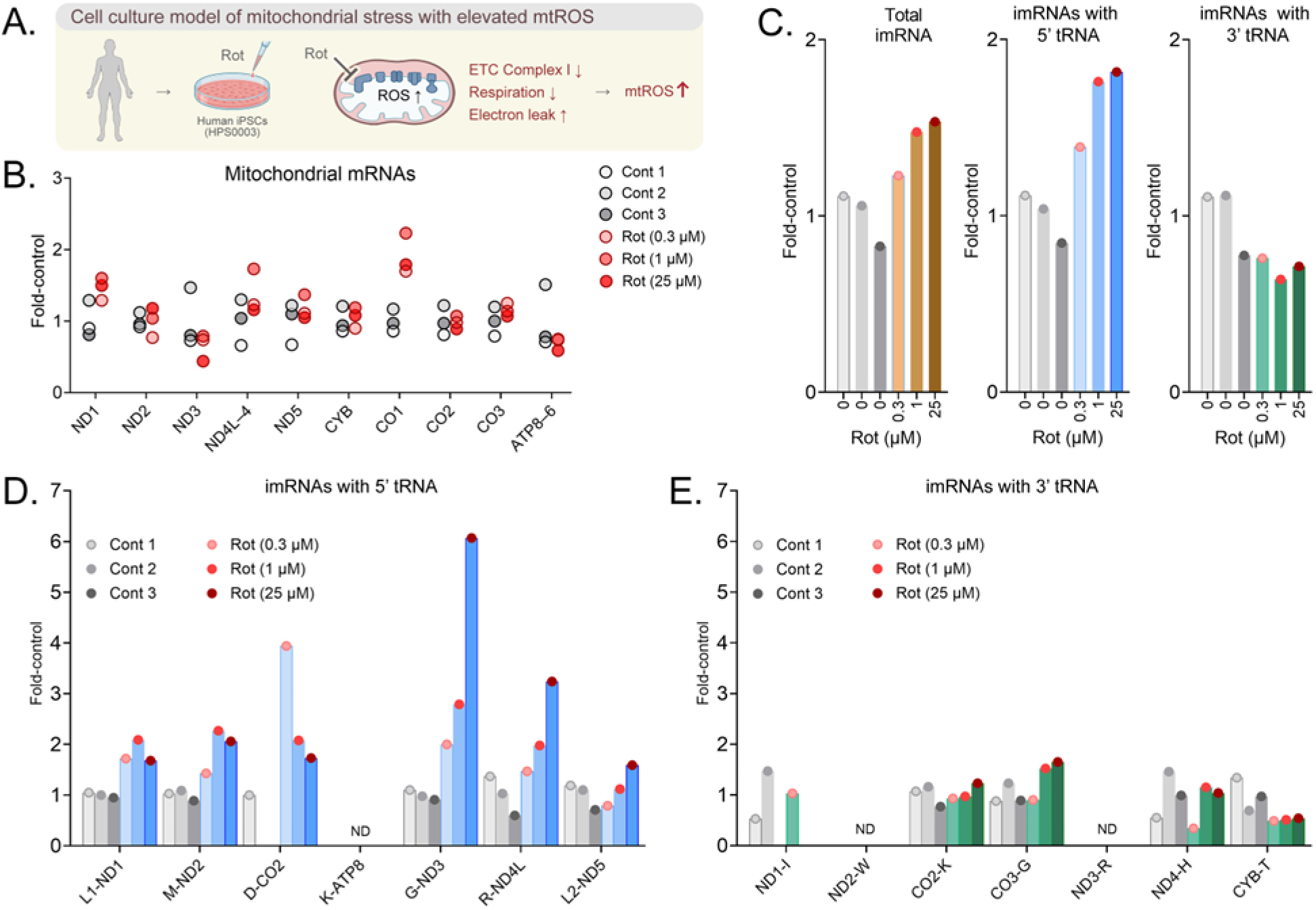
Mitochondrial mRNA levels and imRNA analysis in rotenone-treated human iPSCs. **A.** Human induced pluripotent stem cells (iPSCs) treated with the ETC complex I inhibitor rotenone provide a cell culture model of mitochondrial stress with elevated mtROS. **B.** Mitochondrial mRNA levels in iPSCs treated with increasing doses of dimethyl sulfoxide (Cont 1–3; dots in shades of grey) and rotenone (Rot; shades of red). For brevity, mitochondrial mRNAs were labeled on the x-axis by shortened gene symbols without the “MT-” prefix. *ND4L–4*, bicistronic transcript encoding *MT-ND4L* and *MT-ND4*; *ATP8–6*, bicistronic transcript containing *MT-ATP8* and *MT-ATP6*. **C.** Total imRNA abundance and cumulative abundance of imRNAs retaining 5’ or 3’ tRNAs. **D–E.** Estimated levels of individual imRNAs with 5’ tRNA (panel D) and those with 3’ tRNA (panel E). Individual imRNAs are labeled on the x-axes with the uncleaved junction identifiers (UJIs) they contain; for instance, L1–ND1 marks an ND1 imRNA retaining a 5’ tRNA L1 extension (Supplementary Table 2). In all graphs, values were normalized to the control mean and expressed as multiples of control mean (fold-control). Bars show individual cultures with distinct treatments. Statistical analysis was not performed as each condition is represented by a single biological replicate. DMSO, dimethyl sulfoxide; imRNA, immature mRNA; iPSCs, induced pluripotent stem cells; ND, not detected; Rot, rotenone.

### Specific detection and quantification of imRNAs despite nuclear-encoded mtDNA segments

Although short reads generally cannot establish the complete structure of their source molecules, reads spanning an uncleaved gene boundary provide direct evidence that the corresponding sequences remain unprocessed. Short reads, however, are prone to nonspecific mapping to multiple transcriptomic regions due to sequence similarity. The documented presence of nuclear-encoded mitochondrial segments (NUMTs) with partial or exact sequence matches to mtDNA was particularly challenging in our approach, requiring additional measures to minimize nonspecific detection of nuclear transcripts while identifying imRNAs in mtDNA-derived transcripts. We addressed this in three steps. (i) Sequence comparisons between the mitochondrial and nuclear genomes guided our UJI selection. (ii) Analysis of UJI-containing NUMT expression patterns in six mouse tissues suggested that these critical nuclear regions might be transcriptionally inactive. (iii) Finally, the absence of sequence matches for selected UJIs in the mouse and human ρ^0^ datasets, accompanied by successful detection of all UJIs in control cells with intact mtDNA (**fig 4G**), supported specific detection of mitochondrial transcripts without contamination from the nuclear transcriptome. To estimate relative imRNA abundance, we normalized levels of UJIs not to a predefined common factor, such as total transcript abundance, but to levels of nearby segments within the cognate mRNA, a local denominator specific to each UJI. This normalization helps distinguish changes in the representation of processing intermediates from changes in overall mitochondrial RNA abundance or individual transcript levels. The efficacy of this approach was supported by imRNA results obtained with two datasets representing opposite extremes in mitochondrial transcript abundance: *Mterf4*-mKO with mtRNA accumulation and *Tfam*-mKO with severe mtRNA deficiency (**fig 6A, B**).

### Applicability and potential for rapid adoption

Hardware requirements, simplicity, and accessibility of a new bioinformatics tool, in addition to its promise, are critical determinants of its potential value in research. The imRNA tool presented here can readily be applied to available raw RNA-seq datasets stored in a compressed format as fastq.gz files, without decompression or preprocessing. The algorithm processes read files in parallel across multiple CPU cores to generate an imRNA report in a few hours on a standard personal computer. We anticipate that these features will encourage its adoption by researchers, facilitating research in mitochondrial RNA biology and potentially enabling the discovery of disease-specific imRNA signatures. The imRNA data extracted from RNA-seq data might ultimately serve as additional readouts in basic research and add a new dimension to diagnostic markers in clinical practice.

### An imRNA signature as a potential indicator of oxidative stress and testable predictions for future studies

The application of imRNA analysis to mouse models with a range of mitochondrial perturbations, including elevated mtROS, revealed significant insights into the regulation of mitochondrial gene expression under oxidative stress. The comparisons among these models supported an association between oxidative stress and selective elevation of imRNAs with 5’ tRNA extensions. Mitochondrial defects in translation, transcription, or mtDNA replication did not reproduce the imRNA pattern observed in SOD2-deficient hearts. Furthermore, ANT1 overexpression, a model of mitochondrial import stress associated with reduced mitochondrial hydrogen peroxide production, was accompanied by decreased rather than increased levels of imRNAs with 5’ tRNA. Together, these observations argue against the pattern being an inevitable consequence of respiratory dysfunction, altered mitochondrial transcript abundance, or mitochondrial stress in general. The qualitative reproduction of this pattern in rotenone-treated human cells extends the association across species. It should be emphasized that, although they suggest a potential association, our findings do not yet establish the imRNA pattern as a specific biomarker of elevated mtROS. Establishing specificity will require experiments that measure imRNA levels and mitochondrial redox status in the same samples, manipulate ROS independently of respiratory dysfunction, and test whether reducing oxidative stress reverses the imRNA response. Analysis of *Mterf4*-mKO samples (**fig 6A**), suggesting no increase in imRNA levels despite overall mtRNA accumulation and severely compromised mitochondrial function, might provide an initial lead. Given the stability of imRNA levels in *Mterf4*-mKO mice, our model predicts an absence of mtROS elevation, a testable hypothesis for future investigation. A discordant result would instead help define the conditions under which the signature arises.

### Sequential cleavage model of mitochondrial tRNA maturation does not apply to mRNA maturation

Apart from imRNA analysis under mitochondrial stress, our study provides valuable observations on a particular feature of mitochondrial RNA molecules under physiological conditions, with immediate implications for widely used methods of gene-level expression analysis. Direct RNA sequencing revealed that healthy mouse hearts contain polyadenylated mitochondrial transcripts with complete mRNA coding sequences and retained tRNA extensions, including molecules spanning multiple uncleaved junctions (**fig 3**). These findings indicate that mRNA 3’-end formation and polyadenylation can precede cleavage at upstream junctions and do not require prior release of an individual mRNA from its precursor. Thus, mitochondrial RNA maturation does not follow a single obligatory sequence across the polycistronic transcript. This flexibility is compatible with ordered processing at an individual tRNA, where RNase P-mediated 5’ cleavage generally precedes RNase Z-mediated 3’ cleavage (Reinhard, Sridhara, & Hällberg, 2017). Although our observations are consistent with this local processing order, they do not establish direct coupling between 5’-end cleavage of an mRNA and its polyadenylation. Our findings indicate a clear distinction between tRNA and mRNA maturation at the level of individual molecules: while 5’-end processing of a tRNA molecule precedes its 3’-end processing, the two ends of an mRNA molecule can be processed or remain uncleaved independently of each other. These observations on mitochondrial mRNA maturation might have significant implications for current practices of RNA-seq gene expression analysis as discussed below.

### Potential overestimation of mitochondrial mRNA levels in typical RNA-seq expression analyses

We observed, through direct RNA sequencing, that a significant proportion (24–34%) of mRNA-containing polyadenylated transcripts retained uncleaved sequences in healthy mouse hearts (**fig 3B**). This ratio describes the sequenced polyadenylated population rather than an unbiased estimate of all cellular mitochondrial transcripts. Nevertheless, it shows that polyadenylate (polyA) selection enriches a mixture of mature and incompletely processed mitochondrial RNAs. This finding, together with previous direct RNA-sequencing studies in models of perturbed mitochondrial biology, establishes a critical property of mitochondrial RNA maturation, warranting additional consideration when estimating mRNA levels using RNA-seq. In nuclear pre-mRNA maturation, polyadenylation is performed only after both 5’-end and 3’-end processing are complete. Accordingly, selecting polyA-containing RNA is largely equivalent to obtaining mature mRNA. Mitochondrial mRNA-containing transcripts, however, despite carrying polyA tails, can be maturation intermediates that are not ready for translation. As a result, polyA selection captures both mature mitochondrial mRNAs and polyadenylated processing intermediates containing tRNA or other mitochondrial sequences. Consequently, polyA selection alone does not distinguish mature mitochondrial mRNAs from incompletely processed transcripts. Based on this consideration, we argue that current methodologies for transcriptomic analyses might not be ideal for estimating mitochondrial gene expression. While polyA-mediated RNA selection is practical for obtaining nuclear mRNAs for downstream assays, this approach might result in an overestimation of mitochondrial mRNA abundance in downstream gene-level expression analyses.

### Limitations and mechanistic implications for further investigation

This study has an important limitation that warrants cautious interpretation. The suggested relationship between mtROS and imRNA levels is both unidirectional and indirect. Therefore, it should not be interpreted as evidence that imRNAs themselves are either detrimental or protective. Our findings indicate that SOD2 deficiency and experimentally elevated mtROS can increase imRNAs with 5’ tRNAs, but they do not establish whether these species are byproducts of mitochondrial damage or components of an adaptive response that promotes survival under conditions of supraphysiological ROS. Any mechanistic interpretation beyond this point remains speculative; however, a particularly attractive possibility may be worth considering.

A potential mechanism that might underlie imRNA elevation is inhibition of mitochondrial mRNA maturation by mtROS through targeting of polycistronic RNA processing. Such inhibition could decrease mitochondrial translation and consequently limit electron transport chain activity, creating a negative-feedback loop that restrains further mtROS production. Once mtROS levels return to baseline, subsequent processing could rapidly restore the availability of mature transcripts and support protein synthesis to rebuild the electron transport chain. Importantly, this mechanism could constitute a nucleus-independent homeostatic circuit operating autonomously within individual mitochondria to modulate local ROS production, consistent with the long-proposed model of redox-mediated control of mitochondrial gene expression (John F. Allen, 1993; J. F. Allen, 2015), but acting at the level of RNA maturation and translation rather than transcription. Such a model would imply that eukaryotic cells could contain heterogeneous mitochondrial subpopulations with distinct rates of ROS production and protein synthesis. Recent studies investigating intracellular mitochondrial heterogeneity (Ryu et al., 2024), an emerging area of research facilitated by advances in high-resolution and single-organelle approaches, support the plausibility of this concept.

Nevertheless, experimental evaluation of this model presents significant technical challenges, particularly because it requires quantitative measurements at the level of individual mitochondria. Opposing changes among mitochondrial subpopulations can be obscured in single-cell or tissue-level analyses. This limitation is conceptually similar to that of bulk RNA sequencing, where changes in individual cells can remain undetected, while single-cell RNA-seq may help identify them (Stegle, Teichmann, & Marioni, 2015). Accordingly, a valuable next step would be to develop, when technologically feasible, a high-throughput method for organelle-level transcriptome analysis, which could be termed single-mitochondrion RNA sequencing (smtRNA-seq), along with mitochondrially targeted stable reporters of prior ROS exposure. These approaches could enable measurements of imRNA abundance and ROS exposure at the single-mitochondrion level. The question of whether the proposed ROS-sensitive regulatory mechanism operates within mitochondria is currently under investigation.

## 5. Acknowledgements

We thank Jop van Berlo and Yuk Sham from the University of Minnesota for their valuable suggestions and support. Members of the Julia Liu and Elizabeth Murphy labs provided valuable input through critical discussions. We also thank Murat Nazlimoglu for technical advice and guidance on computational tools.

## 8. Supplementary figure

**Figure S1.**
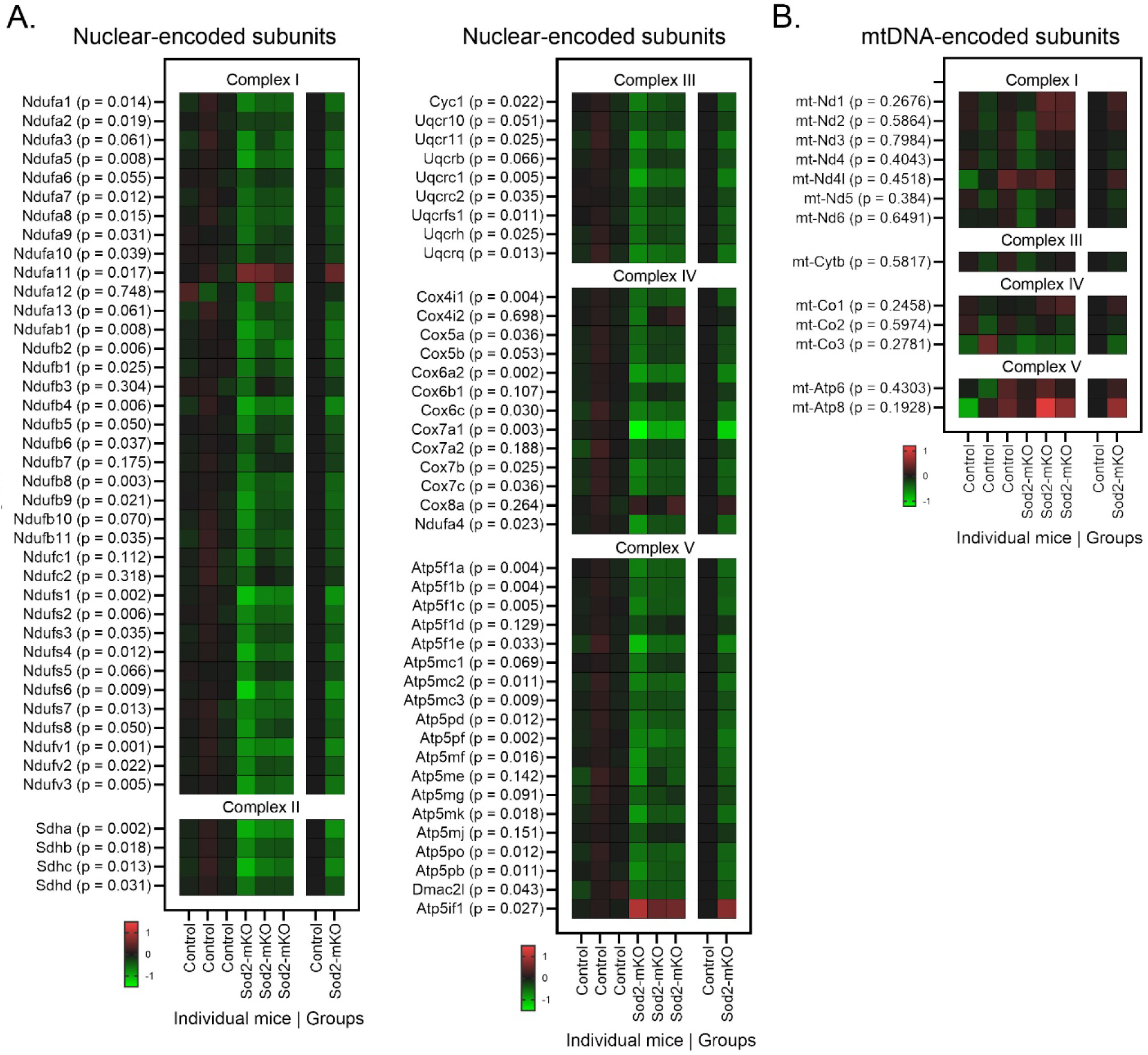
Respiratory chain subunit expression in SOD2-deficient hearts. Nuclear-encoded and mtDNA-encoded respiratory chain mRNA levels in heart tissues from control and Sod2-mKO mice were estimated through typical gene-level expression analysis of RNA-seq data. All known structural subunits expressed in heart tissues were analyzed and reported, with no data exclusion. **A–B.** Heatmaps of nuclear-encoded **(A)** and mtDNA-encoded **(B)** subunit expression. The mRNAs are grouped by ETC complex and labeled with official gene symbols, with p values provided in parentheses. Expression levels are reported as log_2_^FC^, where FC is fold-change relative to the control-group average. Green squares indicate higher while red represents lower expression levels in SOD2-deficient hearts compared to the controls. Each column above “Individual mice” represents a single mouse; those above “Groups” represent the average values for three mice in each group. Sod2, superoxide dismutase 2; mKO, muscle-specific knockout; mtDNA, mitochondrial DNA.

## 9. Supplementary tables

**Supplementary Table 1.**
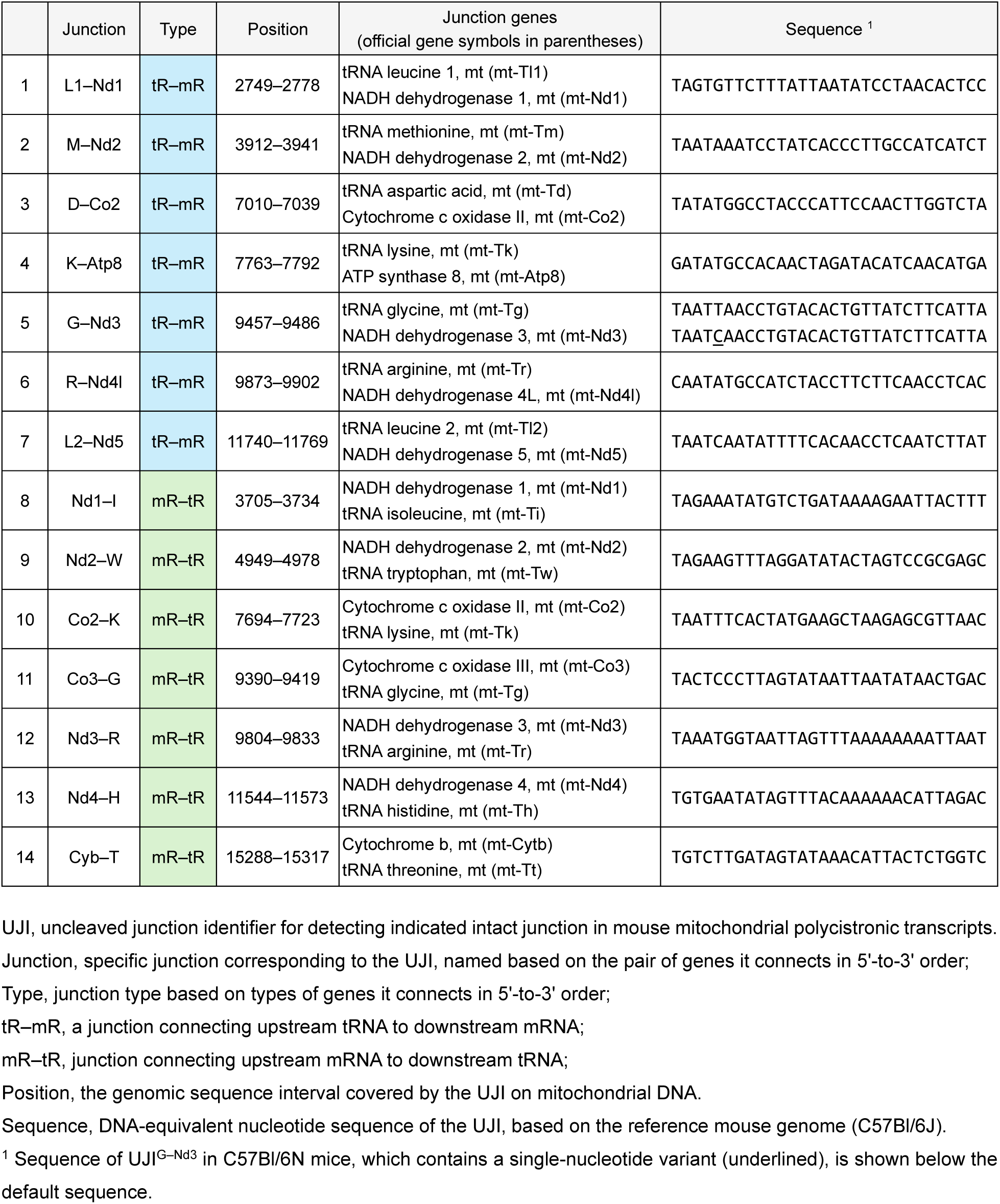
Nucleotide sequences of mouse UJIs.

**Supplementary Table 2.**
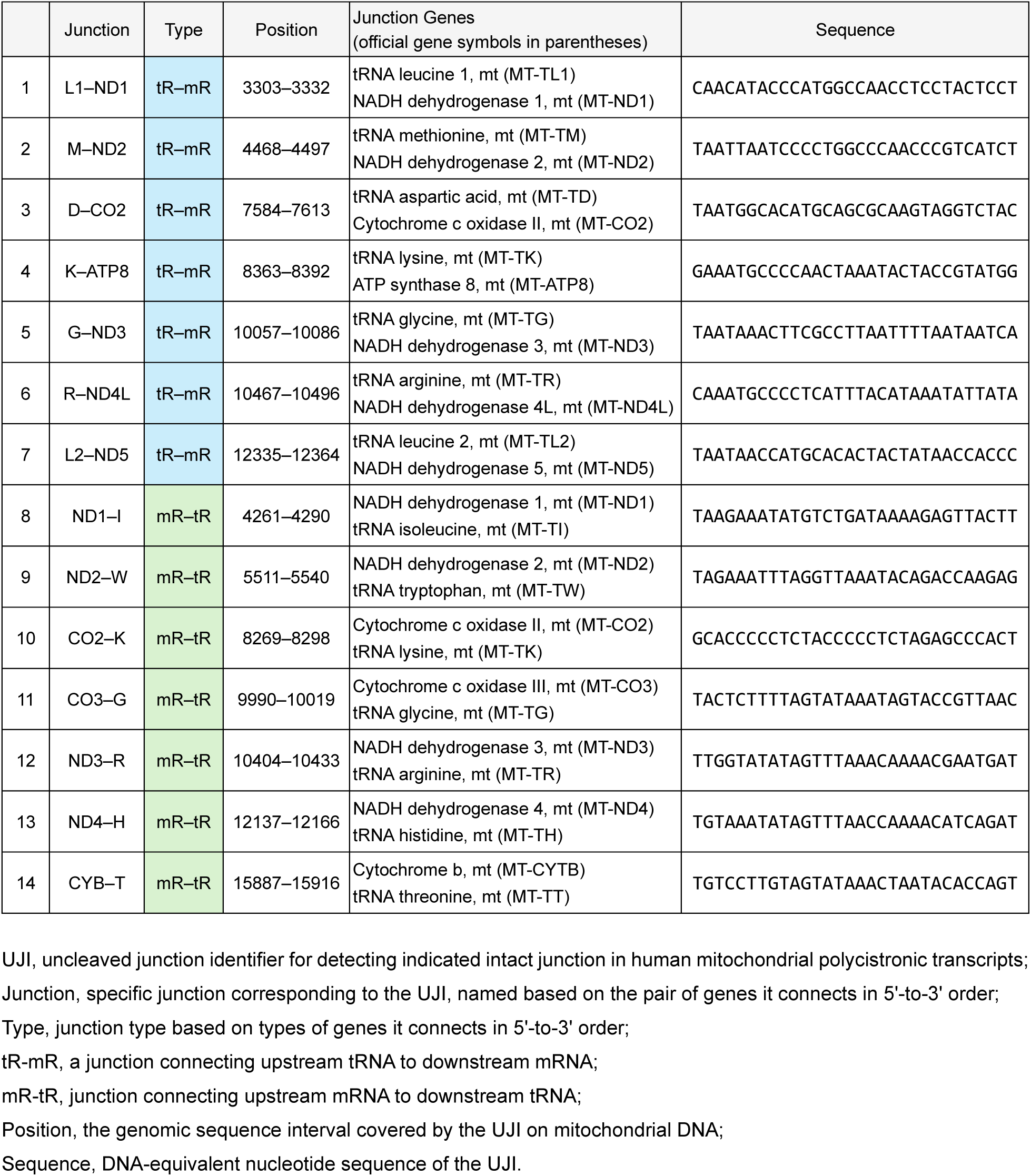
Nucleotide sequences of human UJIs.

**Supplementary Table 3.** Specific detection of mitochondrial transcripts using mouse UJIs in mouse embryonic fibroblasts and mtDNA-depleted cells derived from them (dataset #SRP134850).

| UJI | Junction | Type | Ct #1 | Ct #2 | Ct #3 | ρ<sup>0</sup> #1 | ρ<sup>0</sup> #2 | ρ<sup>0</sup> #3 |
|---|---|---|---|---|---|---|---|---|
| 1 | L1–Nd1 | tR–mR | 6531 | 5542 | 7976 | 0 | 0 | 0 |
| 2 | M–Nd2 | tR–mR | 536 | 492 | 610 | 0 | 0 | 0 |
| 3 | D–Co2 | tR–mR | 28 | 28 | 51 | 0 | 0 | 0 |
| 4 | K–Atp8 | tR–mR | 102 | 145 | 287 | 0 | 0 | 0 |
| 5 | G–Nd3 | tR–mR | 46 | 73 | 95 | 0 | 0 | 0 |
| 6 | R–Nd4l | tR–mR | 52 | 76 | 109 | 0 | 0 | 0 |
| 7 | L2–Nd5 | tR–mR | 21 | 40 | 36 | 0 | 0 | 0 |
| 8 | Nd1–I | mR–tR | 116 | 166 | 305 | 0 | 0 | 0 |
| 9 | Nd2–W | mR–tR | 605 | 558 | 773 | 0 | 0 | 0 |
| 10 | Co2–K | mR–tR | 71 | 117 | 235 | 0 | 0 | 0 |
| 11 | Co3–G | mR–tR | 41 | 53 | 91 | 0 | 0 | 0 |
| 12 | Nd3–R | mR–tR | 28 | 47 | 71 | 0 | 0 | 0 |
| 13 | Nd4–H | mR–tR | 16 | 27 | 22 | 0 | 0 | 0 |
| 14 | Cyb–T | mR–tR | 119 | 126 | 231 | 0 | 0 | 0 |

UJI, uncleaved junction identifier for detecting indicated intact junction in mouse mitochondrial polycistronic transcripts.
Junction, specific junction corresponding to the UJI, named based on the pair of genes it connects in 5'-to-3' order;
Type, junction type based on types of genes it connects in 5'-to-3' order;
tR-mR, a junction connecting upstream tRNA to downstream mRNA;
mR-tR, junction connecting upstream mRNA to downstream tRNA;
Ct 1–3, UJI counts in sequencing reads from three biological replicates of mouse embryonic fibroblasts (MEFs) with intact mtDNA (control samples);
ρ<sup>0</sup> 1–3, UJI counts in sequencing reads from three biological replicates of MEFs devoid of mtDNA (ρ<sup>0</sup> samples);
See Supplementary Table 1 for mouse UJI nucleotide sequences and Methods for details on RNA-seq data.

**Supplementary Table 4.** Specific detection of mitochondrial transcripts using human UJIs in RNA-seq data from human colon tumor cells line 116 and mtDNA-depleted cells derived from them (dataset #SRP093357).

| UJI | Junction | Type | Ct #1 | Ct #2 | Ct #3 | ρ<sup>0</sup> #1 | ρ<sup>0</sup> #2 | ρ<sup>0</sup> #3 |
|---|---|---|---|---|---|---|---|---|
| 1 | L1–ND1 | tR–mR | 104 | 110 | 100 | 0 | 0 | 0 |
| 2 | M–ND2 | tR–mR | 96 | 91 | 106 | 0 | 0 | 0 |
| 3 | D–CO2 | tR–mR | 61 | 55 | 43 | 0 | 0 | 0 |
| 4 | K–ATP8 | tR–mR | 82 | 68 | 84 | 0 | 0 | 0 |
| 5 | G–ND3 | tR–mR | 594 | 615 | 525 | 0 | 0 | 0 |
| 6 | R–ND4L | tR–mR | 224 | 206 | 235 | 0 | 0 | 0 |
| 7 | L2–ND5 | tR–mR | 327 | 317 | 278 | 0 | 0 | 0 |
| 8 | ND1–I | mR–tR | 173 | 140 | 146 | 0 | 0 | 0 |
| 9 | ND2–W | mR–tR | 142 | 145 | 150 | 0 | 0 | 0 |
| 10 | CO2–K | mR–tR | 49 | 47 | 47 | 0 | 0 | 0 |
| 11 | CO3–G | mR–tR | 463 | 465 | 449 | 0 | 0 | 0 |
| 12 | ND3–R | mR–tR | 383 | 339 | 308 | 0 | 0 | 0 |
| 13 | ND4–H | mR–tR | 708 | 638 | 510 | 0 | 0 | 0 |
| 14 | CYB–T | mR–tR | 170 | 130 | 133 | 0 | 0 | 0 |

UJI, uncleaved junction identifier for detecting indicated intact junction in human mitochondrial polycistronic transcripts.
Junction, specific junction corresponding to the UJI, named based on the pair of genes it connects in 5'-to-3' order;
Type, junction type based on types of genes it connects in 5'-to-3' order;
tR-mR, a junction connecting upstream tRNA to downstream mRNA;
mR-tR, junction connecting upstream mRNA to downstream tRNA;
Ct 1–3, UJI counts in sequencing reads from three biological replicates of human colon tumor (HCT) cells with intact mtDNA (control samples);
ρ<sup>0</sup> 1–3, UJI counts in sequencing reads from three biological replicates of HCT cells devoid of mtDNA (ρ<sup>0</sup> samples);
See Supplementary Table 2 for human UJI nucleotide sequences and Methods for details on RNA-seq data.

**Supplementary Table 5.** Publicly available mouse RNA-seq datasets utilized in this study.

| Sample | Data identifier | Biological source of RNA | Dataset identifier | Generated by |
|---|---|---|---|---|
| SOD2 control heart #1 | ERR2356265 | Heart tissue samples from *Sod2<sup>fl/fl</sup>* mice | NCBI BioProject, PRJEB25258 | Larsson group (Kauppila et al., 2018) |
| SOD2 control heart #2 | ERR2356266 |  |  |  |
| SOD2 control heart #3 | ERR2356267 |  |  |  |
| SOD2-deficient heart #1 | ERR2356274 | Heart tissue samples from *Sod2<sup>fl/fl</sup>*;*Ckmm-cre* mice |  |  |
| SOD2-deficient heart #2 | ERR2356275<br>ERR2356276 |  |  |  |
| SOD2-deficient heart #3 | ERR2356277 |  |  |  |
| Ant1 control muscle #1 | SRR31395272 | Quadricep skeletal muscle from wild-type mice | NCBI BioProject, PRJNA1188003; SRA, SRP546145 | Chen group (Brennan et al., 2026) |
| Ant1 control muscle #2 | SRR31395273 |  |  |  |
| Ant1 control muscle #3 | SRR31395274 |  |  |  |
| Ant1 control muscle #4 | SRR31395275 |  |  |  |
| Ant1-Tg muscle #1 | SRR31395267 | Quadricep skeletal muscle from *Ant1*(*Tg*/+) mice, overexpressing ANT1 |  |  |
| Ant1-Tg muscle #2 | SRR31395269 |  |  |  |
| Ant1-Tg muscle #3 | SRR31395270 |  |  |  |
| Ant1-Tg muscle #4 | SRR31395271 |  |  |  |
| Mterf4 control heart #1 | SRR5337738 | Heart tissue samples from *Mterf4<sup>LoxP/LoxP</sup>* mice | NCBI BioProject, PRJNA378878; SRA, SRP101755 | Larsson group (Kühl et al., 2017) |
| Mterf4 control heart #2 | SRR5337739 |  |  |  |
| Mterf4 control heart #3 | SRR5337740 |  |  |  |
| Mterf4 control heart #4 | SRR5337741 |  |  |  |
| Mterf4 control heart #5 | SRR5337742 |  |  |  |
| Mterf4-mKO heart #1 | SRR5337735 | Heart tissue samples from *Mterf4<sup>LoxP/LoxP</sup>*; +/*Ckmm-cre* mice |  |  |
| Mterf4-mKO heart #2 | SRR5337736 |  |  |  |
| Mterf4-mKO heart #3 | SRR5337737 |  |  |  |
| Tfam control heart #1 | SRR6517698 | Heart tissue samples from *Tfam<sup>LoxP/LoxP</sup>* mice |  |  |
| Tfam control heart #2 | SRR6517699 |  |  |  |
| Tfam control heart #3 | SRR6517700 |  |  |  |
| Tfam-mKO heart #1 | SRR6517693 | Heart tissue samples from *Tfam<sup>LoxP/LoxP</sup>*; +/*Ckmm-cre* mice |  |  |
| Tfam-mKO heart #2 | SRR6517694 |  |  |  |
| Tfam-mKO heart #3 | SRR6517695 |  |  |  |
| Tfam-mKO heart #4 | SRR6517696 |  |  |  |
| Tfam-mKO heart #5 | SRR6517697 |  |  |  |
| Polrmt control heart #1 | SRR5337731 | Heart tissue samples from *Polrmt<sup>LoxP/LoxP</sup>* mice |  |  |
| Polrmt control heart #2 | SRR5337732 |  |  |  |
| Polrmt control heart #3 | SRR5337733 |  |  |  |
| Polrmt control heart #4 | SRR5337734 |  |  |  |
| Polrmt-mKO heart #1 | SRR5337727 | Heart tissue samples from *Polrmt<sup>LoxP/LoxP</sup>*; +/*Ckmm-cre* mice |  |  |
| Polrmt-mKO heart #2 | SRR5337728 |  |  |  |
| Polrmt-mKO heart #3 | SRR5337729 |  |  |  |
| Polrmt-mKO heart #4 | SRR5337730 |  |  |  |

**Supplementary Table 6.** Publicly available cell culture RNA-seq datasets utilized in this study.

| Sample | Data identifier | Biological source of RNA | Dataset identifier | Generated by |
|---|---|---|---|---|
| Control MEFs #1 | SRR6824985 | TM6 MEFs:<br>Mouse embryonic fibroblasts isolated from *Pnpt1<sup>flox/flox</sup>* mice | GEO, GSE111668;<br>SRA, SRP134850 | Teitell group<br>(Shimada et al., 2018) |
| Control MEFs #2 | SRR6824986 |  |  |  |
| Control MEFs #3 | SRR6824987 |  |  |  |
| ρ<sup>0</sup> MEFs #1 | SRR6824988 | TM6 MEFs devoid of mtDNA, generated through ditercalinium dichloride dihydrochloride treatment |  |  |
| ρ<sup>0</sup> MEFs #2 | SRR6824989 |  |  |  |
| ρ<sup>0</sup> MEFs #3 | SRR6824990 |  |  |  |
| Control HCT cells #1 | SRR5020817 | HCT116 human colorectal carcinoma cells | SRA, SRP093357 | Savkovic group<br>(Heller et al., 2017) |
| Control HCT cells #2 | SRR5020814 |  |  |  |
| Control HCT cells #3 | SRR5020816 |  |  |  |
| ρ<sup>0</sup> HCT cells #1 | SRR5020815 | HCT116 cells lacking mtDNA, generated through ethidium bromide treatment |  |  |
| ρ<sup>0</sup> HCT cells #2 | SRR5020813 |  |  |  |
| ρ<sup>0</sup> HCT cells #3 | SRR5020820 |  |  |  |
| Human iPSCs control #1 | SRR31394867 | Human iPSCs treated with 0.001% DMSO | GEO, GSE282273;<br>NCBI BioProject, PRJNA1188018 | Sone group<br>(Hongen et al., 2024) |
| Human iPSCs control #2 | SRR31394868 | Human iPSCs treated with 0.01% DMSO |  |  |
| Human iPSCs control #3 | SRR31394869 | Human iPSCs treated with 0.1% DMSO |  |  |
| Human iPSCs ROT #1 | SRR31394870 | Human iPSCs treated with 0.3 µM rotenone |  |  |
| Human iPSCs ROT #2 | SRR31394871 | Human iPSCs treated with 1 µM rotenone |  |  |
| Human iPSCs ROT #3 | SRR31394872 | Human iPSCs treated with 25 µM rotenone |  |  |

DMSO, dimethyl sulfoxide; ROT, rotenone; iPSCs, induced pluripotent stem cells.

